# Exploring the energetic and conformational properties of the sequence space connecting naturally occurring RNA tetraloop receptor motifs

**DOI:** 10.1101/2024.05.28.596103

**Authors:** John H. Shin, Lena M. Cuevas, Rohit Roy, Steve L. Bonilla, Hashim Al-Hashimi, William J Greenleaf, Daniel Herschlag

## Abstract

Folded RNAs contain tertiary contact motifs whose structures and energetics are conserved across different RNAs. The transferable properties of RNA motifs simplify the RNA folding problem, but measuring energetic and conformational properties of many motifs remains a challenge. Here, we use a high-throughput thermodynamic approach to investigate how sequence changes alter the binding properties of naturally-occurring motifs, the GAAA tetraloop • tetraloop receptor (TLR) interactions. We measured the binding energies and conformational preferences of TLR sequences that span mutational pathways from the canonical 11ntR to two other natural TLRs, the IC3R and Vc2R. While the IC3R and Vc2R share highly similar energetic and conformational properties, the landscapes that map the sequence changes for their conversion from the 11ntR to changes in these properties differ dramatically. Differences in the energetic landscapes stem from the mutations needed to convert the 11ntR to the IC3R and Vc2R rather than a difference in the intrinsic energetic architectures of these TLRs. The conformational landscapes feature several non-native TLR variants with conformational preferences that differ from both the initial and final TLRs; these species represent potential branching points along the multidimensional sequence space to sequences with greater fitness in other RNA contexts with alternative conformational preferences. Our high-throughput, quantitative approach reveals the complex nature of sequence-fitness landscapes and leads to models for their molecular origins. Systematic and quantitative molecular approaches provide critical insights into understanding the evolution of natural RNAs as they traverse complex landscapes in response to selective pressures.

## Introduction

Structured RNA molecules perform critical roles in biology as enzymes, regulators, and scaffolding for ribonuclear protein complexes (Zhang and Doudna 2002; Zappulla and Cech 2004; Vicens and Cech 2006; Strobel and Cochrane 2007; Ponting et al. 2009; Chadee et al. 2010; Halvorsen et al. 2010; Mustoe et al. 2014; Roth et al. 2014; Ganser et al. 2019; Mauger et al. 2019; Phan et al. 2021). There has been considerable focus on predicting RNA tertiary structure, and recent efforts have been fueled by the remarkable success of AlphaFold and related machine-learning approaches in providing accurate three-dimensional models for nearly all folded proteins (Cruz et al. 2012; Jumper et al. 2021; Baek et al. 2021; Lin et al. 2023; Schneider et al. 2023; Das et al. 2023; Abramson et al. 2024). Nevertheless, structure prediction alone is insufficient to describe the biological function of RNAs. RNA function is not directly tied to a single structure, rather it depends on the conformational ensemble sampled by the RNA. There is therefore a critical need for deep, quantitative (and ultimately predictive) understanding of the energetic properties of RNA and its conformational landscape. Fortunately, RNAs have structural and energetic properties that simplify their study and can be used to obtain insights into their fundamental properties.

Structured RNAs consist of highly stable secondary structures and sparse tertiary contacts that simplify their study, unlike their protein counterparts (Brion and Westhof 1997; Batey et al. 1999; Baldwin and Rose 1999; Silverman et al. 1999; Russell et al. 2002; Mathews et al. 2010; Bisaria et al. 2017; Herschlag et al. 2018). RNA molecules tend to fold hierarchically such that the problems of secondary and tertiary structure formation can largely be treated separately (Herschlag 1995; Bisaria et al. 2017; Herschlag et al. 2018). Secondary structure is predominantly governed by the interaction of base pairing residues and their nearest neighbors (Turner et al. 1988; Brion and Westhof 1997; Herschlag et al. 2018; Yesselman et al. 2019b; Shi et al. 2020). Tertiary contacts that connect distal regions in primary and secondary sequence often behave as *motifs*; these motifs exhibit common three-dimensional structures when embedded in multiple complex RNAs (Pley et al. 1994; Costa and Michel 1995; Costa et al.

1997; Batey et al. 1999; Klein et al. 2001; Rozhdestvensky et al. 2003; Ishikawa et al. 2011; Zhang et al. 2011; Mládek et al. 2012; Huang and Lilley 2018; Denny et al. 2018a) (Fig. S1). Further, RNA motifs are energetically modular, as the same energetic effects are observed upon mutation or changing conditions when the same motif is embedded in different RNA scaffolds (Bisaria et al. 2017; Herschlag et al. 2018; Denny et al. 2018a; Bonilla et al. 2021).

We emphasize modularity as it lies at the heart of building generalizable models for complex systems in science and allows insights obtained by studying a particular module to be applied in broader contexts (Hartwell et al. 1999; Csete and Doyle 2002; Hendrix et al. 2005; Guttman and Rinn 2012; Hayden et al. 2015). Modularity in RNA tertiary structure implies that the stability and conformational properties of tertiary motifs can be used to “reconstitute” and predict the properties of the full RNA, precepts that are the foundations of the RNA Reconstitution Model (Bisaria et al. 2017; Herschlag et al. 2018).

Practically, a large number of high-quality measurements are needed to provide information of sufficient depth and breadth to dissect and understand the properties and behavior of RNA motifs. The quantitative analysis of RNA on a massively parallel array developed by Greenleaf and coworkers allows the measurement of thermodynamic parameters for tens of thousands of RNA motifs of interest (Buenrostro et al. 2014; She et al. 2017; Denny et al. 2018b; Yesselman et al. 2019a; Bonilla et al. 2021; Shin et al. 2023). We previously used this method to investigate the energetic and conformational properties of a ubiquitous family of RNA tertiary contact motifs known as GNRA tetraloop • tetraloop receptors (TL • TLRs) (Bonilla et al. 2021; Shin et al. 1. 2023) (Fig. 1). We found that TLRs cluster into classes with common TL specificities and conformational preferences, and we also found that sequence variation within classes weaken the tertiary contact without altering their conformational preferences. In particular, the quintessential 11 nt receptor (Jaeger et al. 1994; Murphy and Cech 1994; Costa and Michel 1995; Cate et al. 1996) (11ntR, Fig. 1A) and its variants preferentially bind to the GAAA TL (Costa and Michel 1997; Ikawa et al. 2001; Geary et al. 2008) whereas the TLRs found in the *V. cholera* c-di-GMP riboswitch (Weinberg et al. 2007; Roth and Breaker 2009; Smith et al. 2010) (Vc2R, Fig. 1B), the group IC3 intron (Ikawa et al. 1999, 2001) (IC3R, Fig. 1C), and their sequence variants form a class of TLRs that do not discriminate between GAAA and other TLs such as GUAA (Ikawa et al. 2001; Geary et al. 2008; Ishikawa et al. 2011; Fiore and Nesbitt 2013). This discrimination appears to stem from an interaction between A8 in the 11ntR and 2ʹA in the GAAA that is missing in the Vc2R and IC3R (Cate et al. 1996; Golden et al. 2005; Smith et al. 2009; Zakrevsky et al. 2021) (Fig. 1). In-depth mutational analyses revealed that the residues within the 11ntR exhibit a mix of cooperative and additive energetic properties, with cooperativity dominating proximal groups and additivity dominating groups that are distal in three-dimensional space in the TLR structure (Shin et al. 2023).

**Figure 1:**
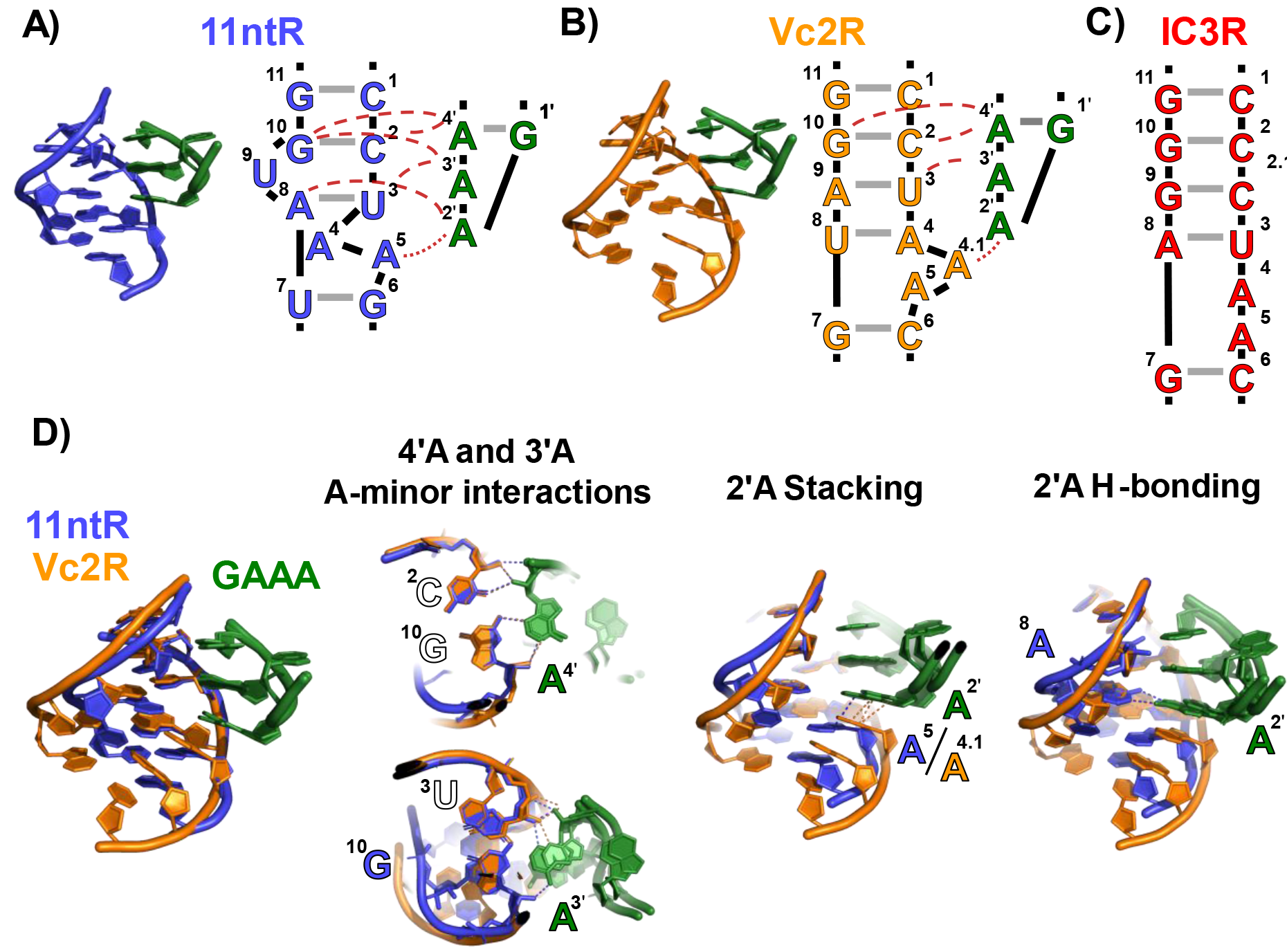
Structured RNA molecules contain modular tertiary contact motifs. (A) Structure and sequence of the 11ntR from the P4-P6 domain of the *Tetrahymena* Group I intron; PDB: 1GID (Cate et al. 1996). Intermolecular hydrogen bonds are denoted by dashed red lines; intermolecular stacking interactions are denoted by dotted red lines. The residues are numbered in accordance with Bonilla *et al*. (Bonilla et al. 2017), and the loop residues of TL are numbered with “primes” to distinguish them from TLR residues. (B) Structure and sequence of Vc2R in the *V. cholera* c-di-GMP riboswitch; PDB: 3MXH (Smith et al. 2010).

The prior quantitative high-throughput biophysical studies extensively described the properties of known TLRs, but of necessity consisted of only a small sampling of the overall TLR sequence space. In particular, the nature of how the properties of TLR variants emerge from or change with sequences remains uninvestigated. Given the immense size of the total TLR sequence space (10^7^ possible 11 or 12 nt sequences) compared to the feasibility limits of the high- throughput assay (10^3^-10^4^ sequences), we chose to explore a subset of the overall sequence space consisting of the mutational pathways from the 11ntR to the IC3R and Vc2R (Fig. 2 A&B). As the IC3R and Vc2R share binding properties (energy and conformation) that differ from the 11ntR, these mutational pathways report on how different TLR binding properties arise from the TLR sequence. Our results suggest that the sequence-energy and sequence-conformation landscapes are not readily parameterized by simple measures such as edit distance from the starting point; rather, specific mutations and sequence intermediates shape the landscapes in complex manners. These idiosyncrasies are interpretable in terms of the underlying molecular interactions in these motifs, highlighting the importance of integrating energetics and molecular models in understanding the sequence landscapes of RNAs, and consequently their function and evolution.

**Figure 2:**
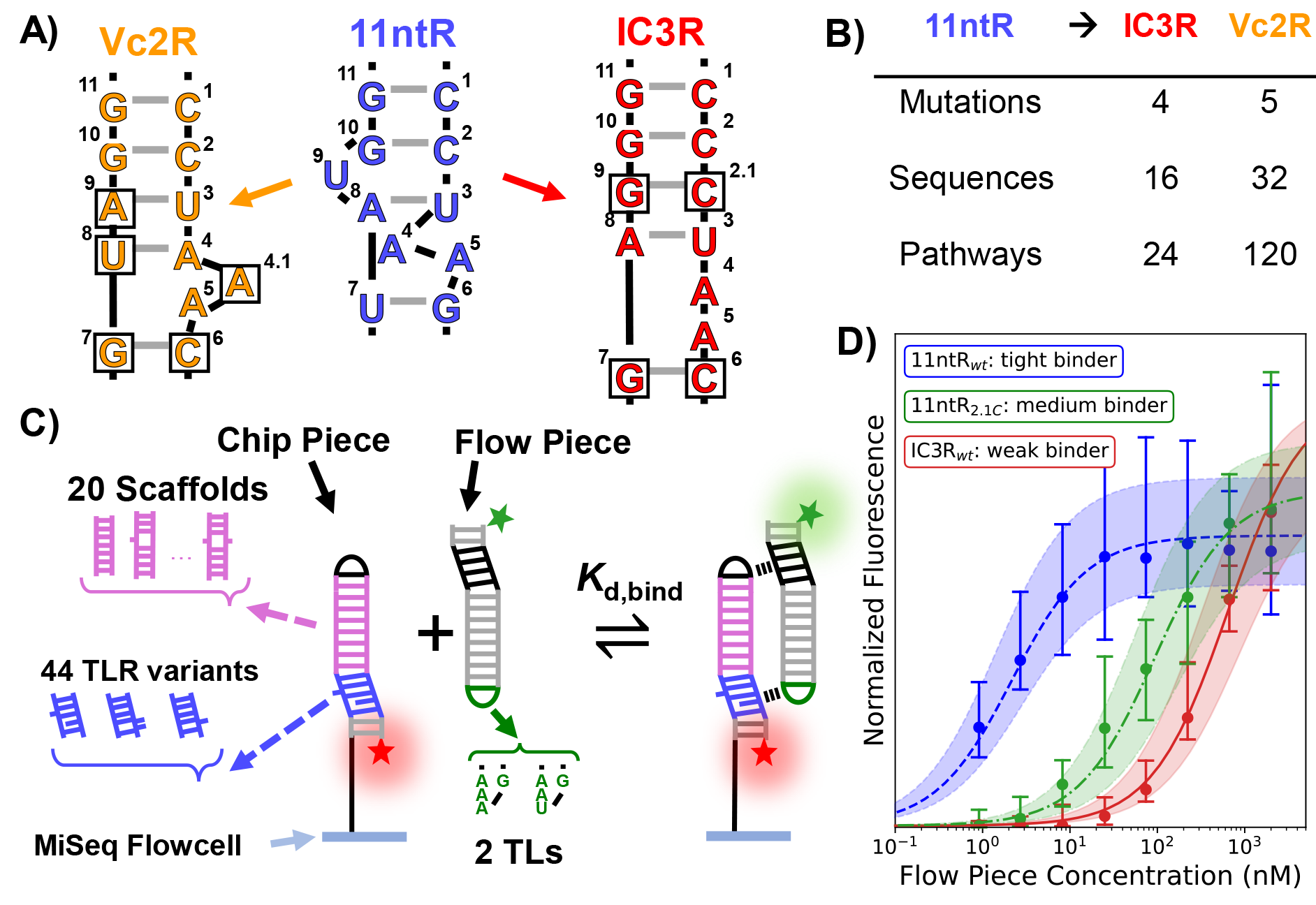
Library design and RNA MaP. (A, B) Conversion from the 11ntR to the IC3R (red) or Vc2R (orange) involves 4 or 5 mutations and 16 or 32 sequence variants, respectively. These variants can be arranged in 24 or 120 single-mutant pathways (Fig. S2). (C) Schematic of the RNA MaP experiment (Buenrostro et al. 2014; She et al. 2017; Denny et al. 2018a). A library of chip piece hairpin sequences is sequenced and transcribed on an Illumina MiSeq flowcell. The chip piece library contains 20 scaffold sequences (pink) and 44 TLR variants (blue) spanning mutational pathways from the 11ntR to the IC3R and Vc2R, resulting in a total of 880 unique sequences. The TLR variants (blue) and a GGAA tetraloop (black) in the chip piece bind to a GAAA or GUAA TL (green) and an R1 TLR (black) in a flow piece hairpin that is flowed onto the MiSeq flowcell. The two tertiary contacts facilitate binding of the flow piece to the immobilized chip piece hairpins. Titrating a fluorescently-labelled flow piece RNA allows for the fitting of isothermal binding isotherms. (D) Example binding isotherms for three TLR sequences: the wt 11ntR (blue; n = 68 clusters), an 11ntR mutant (green; n = 46 clusters) and the wt IC3R (red; n = 78 clusters). Points represent the median fluorescence across all sequence clusters on the flowcell, with error bars depicting the 95% CI obtained via bootstrap analysis. The fit isotherms are depicted as lines going through the data points, with the 95% error interval shaded

## Results

### TLR Anchors and Library Design

The 11ntR, IC3R, and Vc2R (Fig. 1 A-C) were chosen to anchor our exploration of the TLR landscape due to their extensive prior study (Costa and Michel 1995; Ikawa et al. 1999, 2001; Weinberg et al. 2007; Geary et al. 2008; Roth and Breaker 2009; Smith et al. 2010; Bisaria and Herschlag 2015; Bonilla et al. 2017, 2021; Shin et al. 2023). We sampled the landscape spanning these receptors by investigating each stepwise sequence of mutations from the 11ntR to the two other TLRs, focusing on the shortest mutational pathways between the TLR sequences. We substituted each residue of the “left” strand (residues 7-11, Fig. 2A) and, since the “right” strand of the 11ntR (residues 1-6, Fig. 2A) has one fewer nucleotide than the two other receptors, we inserted a residue to this strand in addition to substitutions at other positions. To keep numbering consistent, we denote inserted residues as “X.1” (Fig. 2A). There are four changes between the 11ntR and IC3R that lead to four single, six double, and four triple mutants from the 11ntR in addition to the wild type 11ntR and IC3R. These 16 sequences define 24 single-mutation pathways spanning the TLRs (Fig. 2 A&B, Fig. S2A). Five mutations span the 11ntR and Vc2R receptors, leading to 32 sequences and 120 pathways (Fig. 2 A&B, Fig.S2B). Four of the sequences overlap between the pathways, resulting in 44 unique TLR sequences studied herein.

We utilized the tectoRNA scaffold to study the binding energies and conformational properties of the TLR variants in high throughput on the array platform (Westhof et al. 1996; Jaeger and Leontis 2000; Jaeger et al. 2001; Ishikawa et al. 2013; Denny et al. 2018b; Mitchell et al. 2019; Yesselman et al. 2019a; Bonilla et al. 2021; Shin et al. 2023). The tectoRNA is a heterodimer of RNA hairpins containing two orthogonal TL • TLR pairs that form contacts between the hairpins (Jaeger and Leontis 2000; Jaeger et al. 2001) (Fig. 2C). We inserted our library of TLR variants (Fig. 2C, blue) in the “chip piece” hairpin, where it binds to the TL (Fig. 2C, green) in the complementary “flow piece” hairpin. The main flow piece contained a GAAA TL to bind to the above TLRs, and an alternative flow piece contained a GUAA TL; both flow pieces contained the R1 TLR which forms an orthogonal TL • TLR interaction with the GGAA TL in the chip piece to complete formation of the tectoRNA dimer (Fig. 2C, black) (Jaeger et al. 2001; Mitchell et al. 2019).

The chip piece hairpin contains a variable “scaffold” region between the TLR and TL (Fig. 2C, pink), which enabled us to sample a range of binding geometries between the TLR and its corresponding flow piece TL and obtain information about the TL • TLR conformational landscape. In this study, we used 20 scaffold sequences for a total library of 880 unique chip piece sequence variants (Fig. S3). Binding studies were carried out with the GAAA and GUAA flow pieces in the presence and absence of K^+^; the effects of K^+^ on binding were assessed since the 11ntR contains a K^+^ binding pocket that strengthens the tertiary contact whereas the Vc2R and IC3R do not (Basu et al. 1998; Davis et al. 2007; Lambert et al. 2009; Bonilla et al. 2021; Shin et al. 2023). With the addition of an experimental replicate, we acquired a total of 4400 binding measurements, as described below and in the *Methods*.

### RNA array data collection and analysis

We obtained quantitative binding measurements for our library of TLR variants *via* the previously described quantitative analysis of RNA on a massively parallel array developed by Greenleaf and coworkers (Buenrostro et al. 2014; She et al. 2017; Denny et al. 2018b; Yesselman et al. 2019a; Bonilla et al. 2021; Shin et al. 2023). The 44 TLR x 20 scaffold sequence variants were encoded in a DNA library that was transcribed to RNA on an Illumina MiSeq flowcell (Fig. 2C). The library concentration was controlled to facilitate the generation of multiple “clusters” of chip piece hairpins across the flowcell to serve as technical replicates. We then performed equilibrium tectoRNA binding measurements with the fluorescently-labeled flow piece hairpin to measure binding constants for library construct (Fig. 2D). These experiments were repeated under four different conditions to assay binding of the TLR variants to either a GAAA or GUAA tetraloop in the presence or absence of K^+^. For statistical rigor, we fit isotherms for library variants that had 5 or more replicate clusters on the flowcell as previously described (Denny et al. 2018b) (705-770 of 880 constructs over the experiments, Fig. S4).

The resulting binding free energies (Δ*G* = –*RT* ln *K*eq) were used to calculate ΔΔ*G* values relative to a reference TLR, the wild type 11ntR. The ΔΔ*G* values were largely similar across scaffolds, with a subset of scaffolds having significant different ΔΔ*G* values that reflect different geometric preferences of the TLR (see *Conversion pathways on TLR sequence-conformation landscapes* below). Additionally, several of the scaffolds were destabilizing and resulted in no observable binding for at least a subset of the TLR variants, limiting quantitative comparisons of binding affinities. To improve precision, we report ΔΔ*G* values averaged over a subset of scaffolds that facilitate strong tectoRNA binding (Fig. S7). The effects from conformational changes were separately ascertained by utilizing ΔΔ*G* values for each scaffold.

### Topographies of TLR sequence-energy landscapes

The IC3R and Vc2R share energetic and structural properties that are different from those of the 11ntR: they bind more weakly to the GAAA TL and have broader conformational landscapes (Bonilla et al. 2021). The structure of the GAAA • Vc2R reveals fewer interactions than those found in the GAAA • 11ntR, presumably the origin of its weaker binding and greater flexibility (Fujita et al. 2012) (Fig. 1D). Although no X-ray structure is available for the GAAA • IC3R complex, it has been proposed to make the same base pairs and same interactions as the GAAA • Vc2R (Zakrevsky et al. 2021) (Fig. 1C).

In the simplest case, the similar properties of the Vc2R and IC3R TLRs would translate into similar sequence-energy landscapes spanning from the 11ntR to the IC3R and Vc2R. Such a scenario might involve a steady decrease in binding as the 11ntR is mutated (and interactions are removed) until a point at which additional mutations lead to the formation of new interactions that increase binding in a new energy well. The results described below indicate this simple scenario holds for one of the transition landscapes but not the other. These and additional sequence-binding data combined with structural information give rise to molecular explanations for these differences.

The conversion from the 11ntR to the IC3R follows the simple case of a steady decrease and subsequent increase in binding across the sequence landscape. The 11ntR and IC3R differ by four mutations, with 24 possible pathways for conversion between them (Fig 2 A&B). In every pathway, binding energy is sequentially lost until the weakest intermediate at the third mutation, after which a final mutation leads to tighter binding in the IC3R (Fig. 3A-C, red). We calculated ΔΔ*G* of intermediate TLRs from the previous intermediate (ΔΔ*G*mut) to assess the effect of each mutation at different points along the mutational pathways (Fig. 3E). The first mutation decreases binding by similar amounts, regardless of which mutation is made. At the second step, ΔΔ*G*mut values for each mutation are the same or larger than when it was made first, *e.g.*, the effects of double mutations are additive or compounded to be more deleterious to binding.

**Figure 3.**
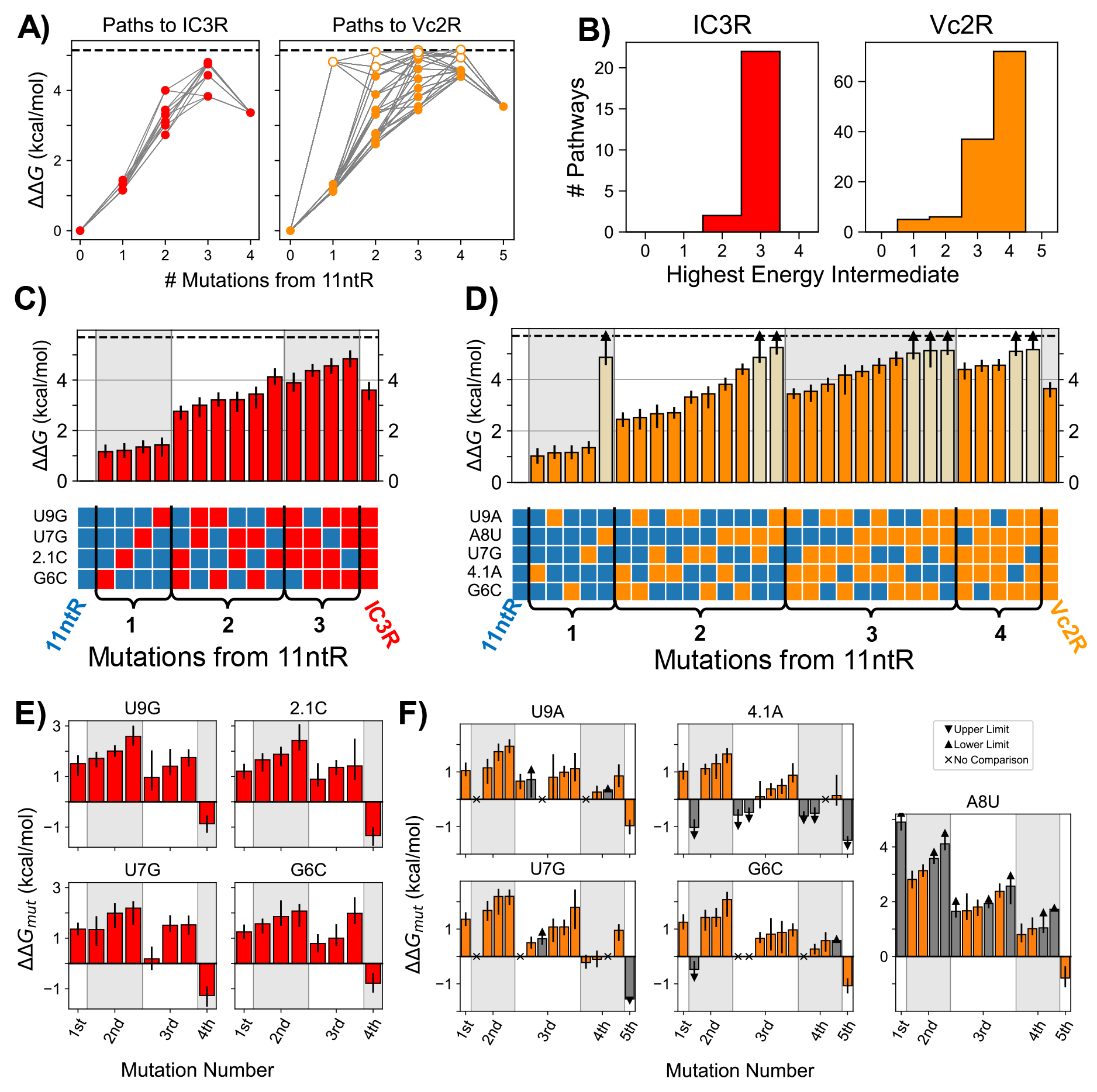
The sequence-energy landscape between naturally occurring TLRs. (A) The sequence-energy landscape for the 11ntR → IC3R conversions (red) and 11ntR → Vc2R conversions (orange) parameterized by the number of mutations from the 11ntR along the x-axis. The y-axis represents ΔΔ*G* relative to the 11ntR (averaged over tectoRNA scaffolds). Each point represents a TLR variant along the conversion landscape, with open circles depicting lower limits for the ΔΔ*G* value. Grey lines connect TLR variants that lie on the same mutational pathway. (B) Distribution of the highest energy intermediate along the mutational pathways from 11ntR to IC3R (left, red) and Vc2R (right, orange). (C, D) The ΔΔ*G* values for TLR sequences along the 11ntR → IC3R pathways (C, red) or the 11ntR → Vc2R pathways (D, orange). ΔΔ*G* values are relative to the 11ntR and averaged over tectoRNA scaffolds; values represent medians of the bootstrapped distributions and error bars represent 95% confidence interval (CI). The specific mutations in each variant are shown in the tables underneath the bar chart, where blue squares represent the original 11ntR sequence and red/orange squares represent mutated sequences towards the IC3R/Vc2R, respectively. Lighter-colored bars represent lower limits for the ΔΔG measurements since the limit of detection (dashed horizontal line) lies within their confidence intervals. (E, F) The mutational effect of each mutation (ΔΔ*G*mut, measured as ΔΔ*G* from the previous mutational intermediate) along the 11ntR → IC3R conversions (E, red) or 11ntR → Vc2R conversions (F, orange); values represent medians of the bootstrapped distributions and error bars represent 95% CI (or limits to the CI). The x-axis is grouped by the order in which the mutation is made, *e.g.*, 1^st^ corresponds to a ΔΔ*G* of a single mutant from the wild type 11ntR. Grey bars correspond to upper or lower limits to ΔΔ*G*mut values (negative or positive values, respectively), and grey “x”s refer to mutations for which neighter quantitative nor qualitative ΔΔ*G*mut values could not be determined.

For the third mutation, the effects are also deleterious (or minimal in one case), but each mutation has a smaller effect than when they were made first or second, suggesting that cooperative interactions are broken or partially broken after the first two mutations. At the last (fourth) step, the mutations that were previously deleterious are now favorable such that all four mutations (whichever is added last) give rise to stronger binding. The shape of this energy landscape suggests that new cooperative interactions are formed in the final IC3R step, but not in earlier steps. Alternatively, the bound TL • TLR complex could consist of two states, an 11ntR-like state and an IC3R-like state, such that mutations create new cooperative interactions in the IC3R-like that are not observed due to stronger binding in the 11ntR-like state until all four mutations are made and IC3R-like binding is favored over 11ntR-like binding.

The conversion landscape from the 11ntR to the Vc2R differs substantially from the conversion pathway to the IC3R described above, as it includes energetically diverse pathways that together define a much less uniform landscape than that for IC3R (Fig. 3A, orange). The conversion requires 5 mutations, leading to 120 possible mutational pathways (Fig. 2A&B). The energetic signatures of these pathways vary, as evidenced by different mutational intermediates representing the highest energy point (Fig. 3B, orange). This variation contrasts with the regular behavior for the IC3R conversion, in which all pathways have the penultimate (third) step as highest in energy (Fig. 3B, red). Despite the differences in the pathways to the VC2R, the final step in each pathway results in stronger binding regardless of the actual mutation made, even for the otherwise highly deleterious A8U mutation (Fig. 3D&F). The increase in binding at the last step––and the different effects of the same mutation at different steps––indicate the cooperative formation of interactions for the Vc2R that enhance TL binding (Fig. 3F), as we also observed for IC3R in the final mutational step (Fig. 3E).

The more featured topography of the 11ntR → Vc2R sequence-energy landscape results from one mutation, A8U, which is much more deleterious than the other mutations when introduced at the first, second or third step (Fig. 3D&F). The distinct energetic effect from mutating residue A8 in the 11ntR presumably arises because it directly hydrogen bonds with 2ʹA in the GAAA. As a result, its removal gives a larger energetic effect than removal of residues that align and aid its interaction (A8 • 2ʹA, Fig. 1D) but do not directly interact with the TL. The role of residue 8 changes in the IC3R and Vc2R (*versus* the 11ntR) and is purported to be the same in both— making a WC pair with residue 4 (Fig. 1B & C). Conversion from the 11ntR to Vc2R requires mutation of A8, eliminating its interaction with 2ʹA before the TLR can adopt the 8 • 4 base pairing architecture (Fig. 1B). In contrast, the retention of A8 in the conversion to IC3R allows it to continue to contribute to binding until too many of the peripheral residues are mutated to support its interaction with 2ʹA. The very different conversion landscapes for IC3R and Vc2R therefore is not a consequence of different physical interactions or different cooperative networks within the final TLRs but rather results from differences in the intermediate species that must be traversed to arrive at each TLR.

Returning to the 11ntR → IC3R landscape, the remarkable smoothness of the pathways might suggest that there are cases where energy landscapes can be modeled as a simple function of sequence space—e.g., Levenshtein distance (Berger et al. 2021)—without considering molecular features. However, two observations indicate that this simplicity is not an intrinsic property of the sequences underlying the 11ntR → IC3R conversion. The first involves the effect of K^+^ which binds to the 11ntR and strengthens its interaction with the GAAA TL (Basu et al. 1998; Davis et al. 2007; Lambert et al. 2009; Bonilla et al. 2021; Shin et al. 2023). While one might expect a simpler landscape upon exclusion of K^+^, the energetic effects of specific mutations instead become more idiosyncratic as residues that had contributed to K^+^ binding no longer do so (Fig. S8A). The second involves binding to a GUAA TL (instead of a GAAA TL), which removes interactions with the 2ʹA (Bonilla et al. 2021), resulting in a more featured landscape (Fig. S8B). Most TLR intermediates bind weakly to the GUAA TL, but some bind with similar or greater affinity than wild type 11ntR, IC3R, or Vc2R. These strongly binding intermediates result in isolated free energy wells, characteristic of a rugged landscape.

### Conversion pathways on TLR sequence-conformation landscapes

TL • TLR interactions stabilize structured RNAs and help specify their conformational state(s). In the previous section, we described changes in the stability of the tertiary interaction as a function of TL • TLR sequence—*i.e.*, the TLR sequence–energy landscape. In this section, we investigated the effects of TLR sequence on the preferred conformation of the TL • TLR interaction (the sequence-conformation landscape). Here, conformation refers to the distance between and orientations of the helices emanating from the TL and TLR, since the formation of the TL • TLR interaction requires these helices in the host RNA molecule to adopt a binding-proficient alignment (Fig 4A). Prior results indicate that the conformational preferences of the IC3R and Vc2R differ from those of the 11ntR (Bonilla et al. 2021). We explored the conformational preferences along the pathways converting the 11ntR to the IC3R or Vc2R, asking where along the pathways the conformational preferences change and where they adopt the preferences of the TLR endpoints.

**Figure 4:**
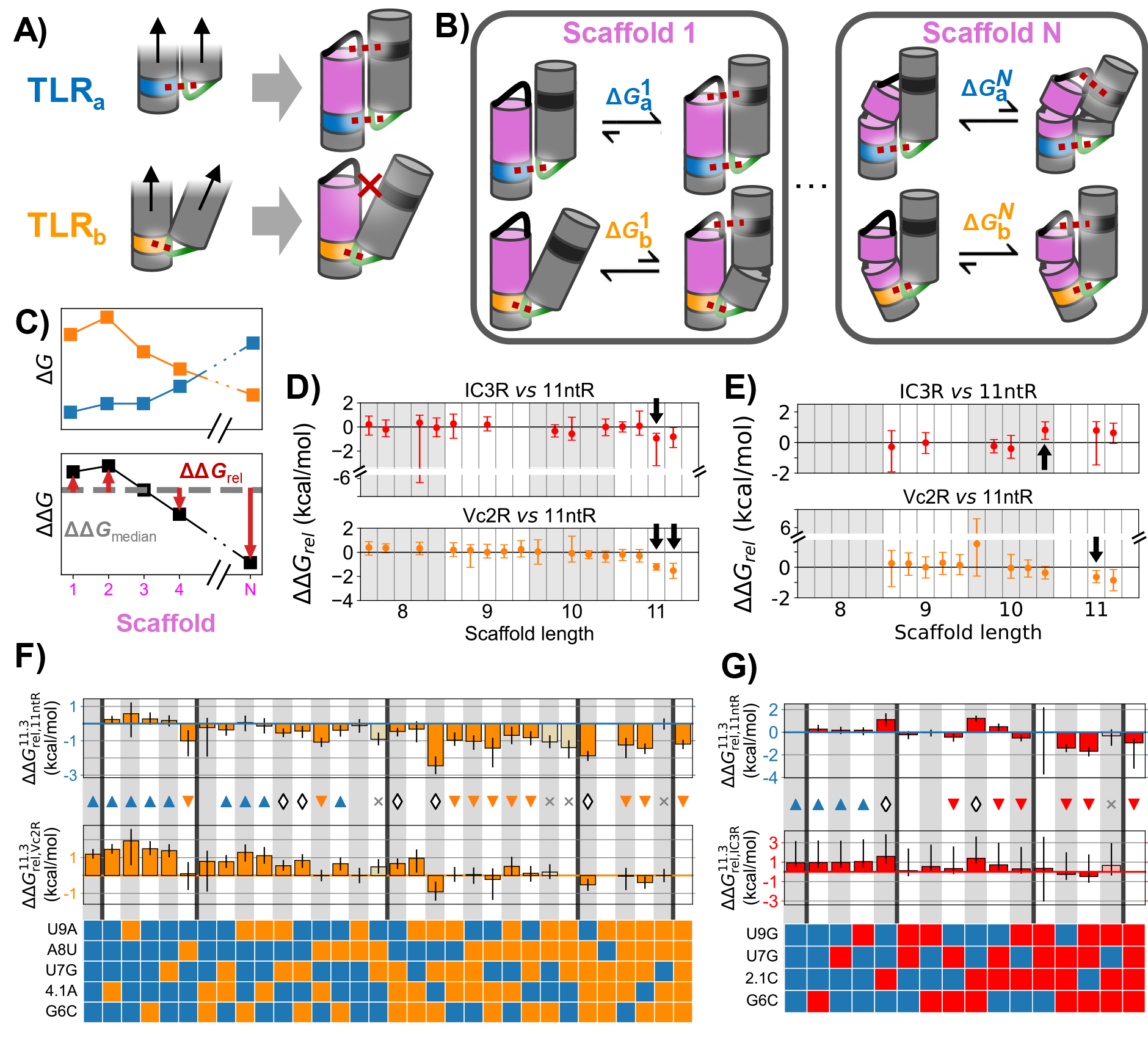
Thermodynamic fingerprints provide information about the sequence-conformation landscape. (A) chematic of how differences in TL • TLR binding geometry and scaffold geometry lead to conformational penalties in a tectoRNA. Two TLRs (a, blue; b, orange) bind to the TL (green) in two orientations. Helical segments are denoted by cylinders and loops/bulges by solid lines connecting them; tertiary contacts are depicted as dashed red lines. The binding of TLRa (blue) aligns the orthogonal TL • TLR pair (black) for binding whereas binding of TLRb (orange) puts the tectoRNA in a conformation that does not favor the orthogonal TL • TLR. (B) Different scaffold geometries can induce tectoRNA conformation changes during binding. The TL • TLRs in (A) are placed into two example tectoRNA scaffolds, one with a straight helix (Scaffold 1) and one with a bulge (Scaffold N). When the geometries of the scaffold and TL • TLR align, tectoRNA interaction is stronger 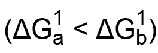; when they do not align, the tectoRNA must change its conformation, resulting in unfavorable binding 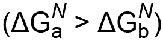. (C) Comparison of thermodynamic fingerprints for two TLRs with different binding geometries. The Δ*G* values (top) across different scaffolds make up their respective thermodynamic fingerprints. ΔΔ*G*rel values (red) are used to compare the thermodynamic fingerprints of different TLR sequences. They are obtained from comparing the ΔΔ*G* between the two sets of Δ*G* values (bottom) to the median ΔΔ*G* value (grey). (D, E) Comparisons of the thermodynamic fingerprints of the 11ntR with the IC3R (red) and Vc2R (orange) binding to a (D) GAAA or (E) GUAA TL. Median and 9 % CIs of ΔΔ*G*rel are depicted across the 20 scaffolds. Scaffolds that induce a significantly different binding conformation (ΔΔ*G*rel ≠ 0) are highlighted by black arrows. (F) The ΔΔ*G*^11.3^ values compared against the 11ntR (top) or IC3R (bottom) for all 11ntR → IC3R variants. Median and 95% CIs for ΔΔ*G*^11.3^ are plotted. Red triangles represent variants with significant differences from the 11ntR, Blue triangles represent variants with significant differences from the IC3R, white diamonds represent variants with significant differences from both. (G) The ΔΔ*G*^11.3^ values compared against the 11ntR (top) or Vc2R (bottom) for all 11ntR → Vc2R variants. Median and 95^th^percentiles for ΔΔ*G*^11.3^are plotted. Orange, blue, and white symbols represent variants with significant differences from the 11ntR, Vc2R, or both, respectively.

The tectoRNA system allows us to compare the conformational preferences of TLRs by varying the scaffold in the chip piece hairpin, shortening/elongating the helix, changing the helical sequence, and/or adding mismatches or internal bulges (Fig 2C, pink, Fig. S3) (Jaeger et al. 2001; Denny et al. 2018a; Yesselman et al. 2019a; Bonilla et al. 2021; Shin et al. 2023). These different scaffolds require conformational adjustments in the tectoRNA to allow the two TL • TLR contacts to form above and below the scaffold (Fig 4B). As a result, the affinity (Δ*G*bind) for each scaffold depends on the energetic cost of adopting distorted, non-optimal tectoRNA conformations (Bonilla et al. 2017; Bisaria et al. 2017; Bonilla et al. 2021; Shin et al. 2023). The set of Δ*G*bind values for a given tertiary interaction as a function of scaffold sequence defines a “thermodynamic fingerprint” for that interaction (Fig 4C, top) (Denny et al. 2018a; Bonilla et al. 2021; Shin et al. 2023). If two TL • TLRs have the same geometric preferences, then they will have identical fingerprints; different fingerprints indicate that the two motifs differ in their binding geometries. To compare the geometric preferences of different TLRs, we calculate ΔΔ*G* values for each scaffold to assess where fingerprints differ (Fig. 4C, bottom). Since each TLR binds the TL with different intrinsic binding strengths, we compare the fingerprints after subtracting the median ΔΔ*G* across the scaffolds for each TLR (ΔΔ*G*rel; Fig. 4C, red arrows). We note that identical fingerprints do not indicate that there are no differences, rather that there are no significant differences in the regions of conformational space explored by the scaffold sequences studied herein.

The thermodynamic fingerprints for the wild type TLRs recapitulate previously described conformational differences between the 11ntR and the IC3R or Vc2R (Bonilla et al. 2021). The longer 11 bp scaffolds generally decrease GAAA binding for the 11ntR more than either the IC3R or Vc2R (Fig. 4D, ΔΔ*G*_rel_ <0, Fig. S9A). This differential effect is decreased or abrogated when binding to the GUAA TL, suggesting that the 11ntR interaction with the 2ʹA that is not present for IC3R or Vc2R is responsible for the different thermodynamic fingerprints and the more narrow fingerprint for the 11ntR than for the IC3R or Vc2R (Fig. 4E, Fig. S9B) (Bonilla et al. 2021).

As 2ʹA makes hydrogen bonds to A8 of the 11ntR but not in the Vc2R or IC3R (Fig. 1), we might expect different conformational landscapes for conversion from the 11ntR to the IC3R *versus* the Vc2R based on whether A8 is mutated, as we saw for the energetic landscapes. To test this model and more broadly investigate the conformational landscape, we compared the thermodynamic fingerprints of intermediate TLRs to their pathway endpoints. In particular, we used the 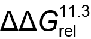 values for Scaffold 11.3 (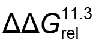 for comparisons to the 11ntR, IC3R, or Vc2R, respectively) as a metric for changes to conformational preferences as it had the most significant difference for the wild type receptors (Fig. 4D, black arrows)

The A8U mutation gives a fingerprint closer to the Vc2R than the 11ntR, as predicted, and most subsequent intermediates along the 11ntR → Vc2R pathways containing A8U maintain a Vc2R- like fingerprint (8 of the 10 sequences with quantitative ΔΔ*G*^11.3^, Fig. 4F). However, one subsequent mutation, G6C, restores the 11ntR conformational preference despite the mutation of A8 (ΔΔ*G*^11.3^ ∼ 0 for A8U/G6C). Residues 6 and 8 are not expected to interact within the receptor, so the non-independent conformational effects may arise indirectly *via* other interactions within the motif.

The other deviating mutant, A8U/G6C/4.1A/U7G, results in a fingerprint that differs from both the 11ntR and Vc2R, indicating that the 11ntR → Vc2R conversion landscape contains potential branching points to alternative TLRs with different binding conformations (Fig. 4F, Fig. S10B). A distinct fingerprint is also seen for the U9A/U7G/G6C triple mutant (Fig. 4F, Fig. S10B). These two intermediates were not outliers in the energetic landscapes, indicating that the conformational and energetic properties of TLR sequences need not track together across sequence space.

The energy and conformational landscapes for the 11ntR → IC3R conversion also differ, even though it does not involve a direct disruption to the A8 • 2ʹA interaction (Fig. 4G). In particular, the 2.1C insertion is associated with a non-11ntR fingerprint (Fig. 4G, Fig. S10A), consistent with the insertion disrupting the interactions between 2C and 3U in the 11ntR with the TL (Fig. 1A&D).

Overall, the sequence-conformation landscapes spanning the conversions between these TLR are complex. Sampling different TLR conformational preferences across sequence space can result in context-dependent energetic penalties and, presumably, additional complexities in adaptive fitness landscapes.

### TL•TLR fitness-stability relationships depend on their RNA hosts

We previously observed that the number of 11ntR sequence variants found in natural RNAs correlates with the stability of its interaction with the TL (Bonilla et al. 2021). This relationship is consistent with a model in which TL • TLR stability is a major driving force for 11ntR selection (Fig. 5A). A strong correlation remains when we include all known natural TLR variants (Fig. 5B). Nevertheless, several sequences occur with much higher frequencies than expected from the correlation, including the IC3R (red point, Fig. 5B). This overrepresentation of the weaker-binding IC3R in natural RNAs introduces the possibility of positive selection for this TLR (and potentially others) based on properties other than tertiary contact stability.

**Figure 5:**
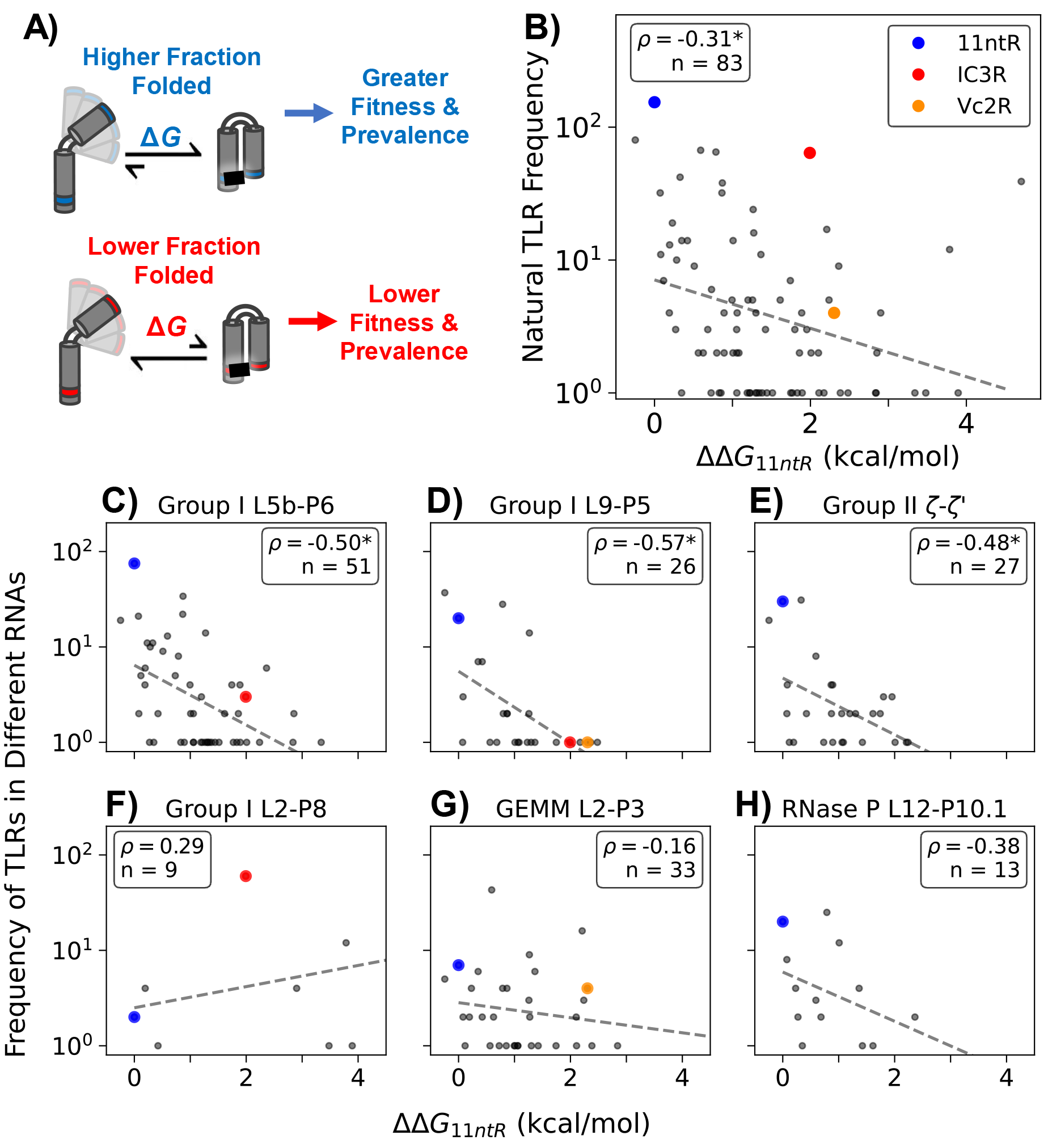
Frequency – stability relationships for natural TLRs. (A) A more stable tertiary interaction could lead to a higher fraction folded of the host RNA, in turn leading to greater fitness and prevalence in biology. (B) The (log) frequency of TLR variants across different host RNAs has a (significantly) negative correlation with the stability of the RNA. (C-H) The frequency-stability correlation is stronger in some host RNA contexts and nonexistent in others. tability is represented as the average ΔΔG from the 11ntR in 30 mM Mg^2+^ taken from Bonilla *et al*. (Bonilla et al. 2021). The three TLRs highlighted in this work are denoted by colored points: blue for the 11ntR, red for the IC3R, and orange for Vc2R. Dashed grey lines plot the log-linear fit between frequency and stability. Significant negative correlations are signified with an asterisk.

To investigate whether TLRs in different RNA contexts might be subject to different selective pressures, we plotted TL • TLR frequency-stability relationships for each of the six RNA locations in which natural TLR variants have been found (Fig. 5B–G). Three of the six maintain the above-noted correlation between frequency and stability, with statistically significant Pearson correlation coefficients of ∼–0.5 (Fig 5C-E, Group I L5b-P6, Group I L9-P5, and Group II ζ-ζʹ). In contrast, two do not have significant correlations (Fig. 5F&G, Group I L2-P8, GEMM L2-P3), and one (RNase P L12-P10.1, Fig. 5H) has too few points and/or too much variation to be statistically significant.

The IC3R is the most represented TLR in the Group I L2-P8 contact (Fig 5F, red), suggesting that this structural context may select for properties other than the higher stability given by the 11ntR. In particular, the TL • TLR with IC3R favors different binding conformations than that with the 11ntR. Nevertheless, the Vc2R, which has a similar binding conformation to the IC3R, is absent from Group I L2-P8 contact, underscoring the possibility of additional selective features as well as limitations from the small number of and biases in the represented sequences from the evolutionary history of these RNAs.

The TLR sequence frequency found L2-P3 of the GEMM riboswitch also does not correlate with stability (Fig. 5G). The need for riboswitches to change structure upon ligand binding may favor TL • TLRs of intermediate strength––strong enough to form but weak enough to be disrupted upon ligand binding (Sudarsan et al. 2008; Smith et al. 2009). Nevertheless, a range of stabilities are present in the GEMM riboswitch TL • TLRs. It will be fascinating to determine if evolution has tuned the stabilities of riboswitches and other RNAs (including the ribosome and spliceosome) to allow the formation of and transition between multiple states and to what extent this tuning differs in different cellular environments.

## Discussion

Evolutionary models often, by necessity, assume simplified adaptive landscapes. Nevertheless, we found that the energy and conformational landscapes spanning the parsimonious conversion pathways from the 11ntR to the IC3R or Vc2R are highly dependent on the intermediate sequences traversed, even for a case where the properties of the endpoints (IC3R and Vc2R) are similar. Comprehensive energetic and conformational data combined with structural models provided explanations for at least some of these behavioral differences. Quantitative multi- dimensional information about these RNA elements––their stabilities and conformational preferences––also allowed us to test models for selection and to provide evidence for the dominance of different selective pressures for the same RNA motifs in different RNA host molecules.

Investigating how the biophysical properties of RNA motifs vary across sequence space is critical for understanding the function and evolution of motifs and their host RNAs. We focused on three naturally occurring TLR motifs in this work, but our systematic quantitative approach is readily adapted to other RNAs. We suggest that this approach will be necessary to establish biochemical understanding, as the biochemical functions of RNAs are enabled by the combined biophysical properties of their elements. Biological understanding can also be revealed from the underlying energetic and conformational landscapes of RNA, which may provide insights into selective driving forces and additional context-dependent constraints. For example, parallel systematic approaches in vitro and in vivo may help dissect the interplay of kinetic and thermodynamic factors for riboswitch regulation in cells and for other functional RNAs.

## Methods

### RNA Array Data Collection

Equilibrium constants for the TLR • TL interactions were derived from fluorescent binding experiments on a modified Illumina MiSeq flowcell as previously described (Buenrostro et al. 2014; She et al. 2017; Denny et al. 2018b). In short, the normalized fluorescence data for molecular replicates (clusters) were resampled in a bootstrap procedure and fit to binding isotherms. We set a minimum number of five clusters per sequence variant to fit equilibrium constants. Four experimental conditions were assayed: TLRs binding to GAAA in 30 mM Mg^2+^ and 30 mM Mg^2+^ + 150 mM K^+^; TLRs binding to GUAA in 30 mM Mg^2+^ and 30 mM Mg^2+^ + 150 mM K^+^. Detailed Materials and Methods are available in the *Supplemental Methods*.

Sequence variants had an average of 41 and 24 clusters for binding to GAAA and GUAA in 30 mM Mg^2+^, and 43 and 43 clusters for binding GAAA and GUAA in 30 mM Mg^2+^ + 150 mM K^+^ (Fig. S4). We identified experimental thresholds for binding based on the fit Δ*G*bind of negative controls in the library, which consist of RNA molecules lacking a tetraloop receptor. We set the cutoff as the 99th percentile of the Δ*G*bind distribution, which differed across experiments (Fig. S5). The Δ*G*bind values for TLR variants were clipped at the cutoff, which should be understood as a lower limit for binding under the respective experimental conditions. The cutoffs were found to be –6.9 kcal/mol and –7.1 kcal/mol for binding to GAAA or GUAA in 30 mM Mg^2+^, and –7.0 kcal/mol and –6.8 kcal/mol for binding to GAAA or GUAA in 30 mM Mg^2+^ + 150 mM K^+^.

Two experiments were performed to measure binding of 11ntR variants to the GAAA tetraloop in 30 mM Mg^2+^; and the replicate data correlated well with a correlation coefficient of 0.94 among non-limit measurements (Fig. S6). The resulting Δ*G*bind distributions were combined by taking the average of the two Δ*G*bind values from each bootstrap (n = 10,000) distribution:

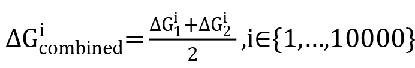

### Statistical analysis

We leveraged the empirical distributions of Δ*G*bind obtained via bootstrapping to directly propagate uncertainty throughout our calculations. ΔΔ*G* values, the difference between variant and wild-type binding, were calculated by subtracting the wild-type Δ*G*bind value from the mutant Δ*G*bind value for each scaffold. Reported Δ*G*bind and ΔΔ*G* values represent the median and 95% confidence interval from the distribution of the average over a subset of 9 scaffold sequences, which were chosen to maximize quantitative (i.e., non-limit) values while omitting scaffolds that induce significantly different conformational penalties (e.g., Scaffolds 11.3 and 11.4, see *Conversion pathways on TLR sequence-conformation landscapes*).

### Thermodynamic fingerprints

We assessed whether the thermodynamic fingerprints of two TLRs differ, as follows. First, we isolated the conformational effect from the binding energy by determining an offset between the Δ*G* values for the TLRs. The following expression was used to calculate this offset:

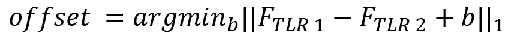

Where 𝐹_𝑇𝐿𝑅 𝑖_ is a vector that contains the thermodynamic fingerprint for TLR *i* (the Δ*G* values for all 20 scaffolds) and || · ||_1_ is the 𝐿^1^ norm, which we chose due to its resilience to outliers (*i.e.*, scaffolds that result in big differences in binding will affect the offset less). From the properties of the 𝐿^1^ norm, we know that the above equation reduces to:

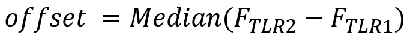

With this offset, we then calculated the ΔΔ*G*rel for a TLR compared to a reference TLR:

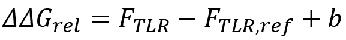

where 𝛥𝛥𝐺_𝑟𝑒𝑙_ represents a vector holding the ΔΔ*G*rel value for all 20 scaffolds. We then calculated each (one-sided) p-value testing for a significantly nonzero ΔΔ*G*rel value:

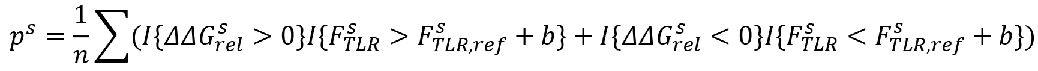

where the superscript 𝑠 represents a calculation for a particular scaffold, 𝐹_𝑇𝐿𝑅_ is the fingerprint of the TLR of interest, and 𝐹_𝑇𝐿𝑅,𝑟𝑒𝑓_ is fingerprint of the reference TLR to which the comparison is made.

The resulting *p*-values were corrected for multiple hypothesis testing using the Benjamini-Hochberg false discovery rate correction (Benjamini and Hochberg 1995) for each of the 11ntR

→ IC3R and 11ntR → Vc2R sequence-conformation landscapes. Significance was assigned if the adjusted p-value is less than 0.05.

### Frequency versus stability fits

Frequency and stability data for 83 naturally-occurring TLR variants were obtained from previously published results (Bonilla et al. 2021). As discussed in the section *TL•TLR fitness- stability relationships depend on their RNA hosts*, we found an empirical log-linear relationship between the relative frequency of a TLR in a given structural context and its relative free energy:

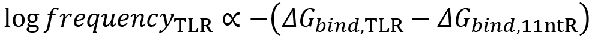

To test this relationship across different RNA host contexts, we calculated the Pearson correlation coefficient between the log frequency and the ΔΔ*G*11ntR for every TLR found in a particular host context. The frequency and ΔΔ*G*11ntR values were resampled *n* = 10,000 times to generate a bootstrapped distribution for the Pearson correlation coefficient. Structural contexts in which the 95% confidence interval for the correlation coefficient did not contain 0 were deemed to have a significant correlation; otherwise, no significant correlation was noted.

## Acknowledgements

We thank members of the Herschlag laboratory for discussion and review of the manuscript. We would like to thank Fanny Sunden for her insights and guidance in designing and running experiments and Joe Yesselman for his computational and conceptual support and pedagogy during library development. This work was supported by the National Institutes of Health (R01 GM132899 to D.H. and H.A.). J.H.S. was supported by the NSF Graduate Research Fellowship Program under Grant DGE-1656518. D.H. serves on the Scientific Advisory Board of Arrakis Therapeutics.

Some of the computing for this project was performed on the Sherlock cluster. We would like to thank Stanford University and the Stanford Research Computing Center for providing computational resources that contributed to these research results. Molecular structures were visualized using PyMOL (Schrödinger, LLC.).

## Notes

### Competing Interest Statement

Daniel Herschlag serves on the Scientific Advisory Board of Arrakis Therapeutics

## References

1. Abramson J, Adler J, Dunger J, Evans R, Green T, Pritzel A, Ronneberger O, Willmore L, Ballard AJ, Bambrick J, et al. 2024. Accurate structure prediction of biomolecular interactions with AlphaFold 3. Nature 1–3.

2. Baek M, DiMaio F, Anishchenko I, Dauparas J, Ovchinnikov S, Lee GR, Wang J, Cong Q, Kinch LN, Schaeffer RD, et al. 2021. Accurate prediction of protein structures and interactions using a three-track neural network. Science 373: 871–876.

3. Baldwin RL, Rose GD. 1999. Is protein folding hierarchic? I. Local structure and peptide folding. Trends Biochem Sci 24: 26–33.

4. Basu S, P. Rambo R, Strauss-Soukup J, H. Cate J, R. Ferré-D’Amaré A, Strobel SA, Doudna JA. 1998. A specific monovalent metal ion integral to the AA platform of the RNA tetraloop receptor. Nat Struct Biol 5: 986–992.

5. Batey RT, Rambo RP, Doudna JA. 1999. Tertiary Motifs in RNA Structure and Folding. Angew Chem Int Ed 38: 2326–2343.

6. Benjamini Y, Hochberg Y. 1995. Controlling the false discovery rate: a practical and powerful approach to multiple testing. J R Stat Soc Ser B Methodol 57: 289–300.

7. Berger B, Waterman MS, Yu YW. 2021. Levenshtein Distance, Sequence Comparison and Biological Database Search. IEEE Trans Inf Theory 67: 3287–3294.

8. Bisaria N, Greenfeld M, Limouse C, Mabuchi H, Herschlag D. 2017. Quantitative tests of a reconstitution model for RNA folding thermodynamics and kinetics. Proc Natl Acad Sci 114: E7688–E7696.

9. Bisaria N, Herschlag D. 2015. Probing the kinetic and thermodynamic consequences of the tetraloop/tetraloop receptor monovalent ion-binding site in P4–P6 RNA by smFRET. Biochem Soc Trans 43: 172–178.

10. Bonilla S, Limouse C, Bisaria N, Gebala M, Mabuchi H, Herschlag D. 2017. Single-Molecule Fluorescence Reveals Commonalities and Distinctions among Natural and in Vitro-Selected RNA Tertiary Motifs in a Multistep Folding Pathway. J Am Chem Soc 139: 18576–18589.

11. Bonilla SL, Denny SK, Shin JH, Alvarez-Buylla A, Greenleaf WJ, Herschlag D. 2021. High-throughput dissection of the thermodynamic and conformational properties of a ubiquitous class of RNA tertiary contact motifs. Proc Natl Acad Sci 118.

12. Brion P, Westhof E. 1997. Hierarchy and Dynamics of RNA Folding. Annu Rev Biophys Biomol Struct 26: 113–137.

13. Buenrostro JD, Araya CL, Chircus LM, Layton CJ, Chang HY, Snyder MP, Greenleaf WJ. 2014. Quantitative analysis of RNA-protein interactions on a massively parallel array reveals biophysical and evolutionary landscapes. Nat Biotechnol 32: 562–568.

14. Cate JH, Gooding AR, Podell E, Zhou K, Golden BL, Kundrot CE, Cech TR, Doudna JA. 1996. Crystal Structure of a Group I Ribozyme Domain: Principles of RNA Packing. Science 273: 1678–1685.

15. Chadee AB, Bhaskaran H, Russell R. 2010. Protein Roles in Group I Intron RNA Folding: The Tyrosyl-tRNA Synthetase CYT-18 Stabilizes the Native State Relative to a Long-Lived Misfolded Structure without Compromising Folding Kinetics. J Mol Biol 395: 656–670.

16. Costa M, Dème E, Jacquier A, Michel F. 1997. Multiple tertiary interactions involving domain II of group II self-splicing introns11Edited by M. Yaniv. J Mol Biol 267: 520–536.

17. Costa M, Michel F. 1995. Frequent use of the same tertiary motif by self-folding RNAs. EMBO J. 14: 1276–1285.

18. Costa M, Michel F. 1997. Rules for RNA recognition of GNRA tetraloops deduced by in vitro selection: comparison with in vivo evolution. EMBO J 16: 3289–3302.

19. Cruz JA, Blanchet M-F, Boniecki M, Bujnicki JM, Chen S-J, Cao S, Das R, Ding F, Dokholyan NV, Flores SC, et al. 2012. RNA-Puzzles: A CASP-like evaluation of RNA three-dimensional structure prediction. RNA 18: 610–625.

20. Csete ME, Doyle JC. 2002. Reverse Engineering of Biological Complexity. Science 295: 1664– 1669.

21. Das R, Kretsch RC, Simpkin AJ, Mulvaney T, Pham P, Rangan R, Bu F, Keegan RM, Topf M, Rigden DJ, et al. 2023. Assessment of three-dimensional RNA structure prediction in CASP15. Proteins Struct Funct Bioinforma 91: 1747–1770.

22. Davis JH, Foster TR, Tonelli M, Butcher SE. 2007. Role of metal ions in the tetraloop–receptor complex as analyzed by NMR. RNA 13: 76–86.

23. Denny SK, Bisaria N, Yesselman JD, Das R, Herschlag D, Greenleaf WJ. 2018a. High-Throughput Investigation of Diverse Junction Elements in RNA Tertiary Folding. Cell 174: 377–390.e20.

24. Denny SK, Bisaria N, Yesselman JD, Das R, Herschlag D, Greenleaf WJ. 2018b. High-Throughput Investigation of Diverse Junction Elements in RNA Tertiary Folding. Cell 174: 377–390.

25. Fiore JL, Nesbitt DJ. 2013. An RNA folding motif: GNRA tetraloop–receptor interactions. Q Rev Biophys 46: 223–264.

26. Fujita Y, Tanaka T, Furuta H, Ikawa Y. 2012. Functional roles of a tetraloop/receptor interacting module in a cyclic di-GMP riboswitch. J Biosci Bioeng 113: 141–145.

27. Ganser LR, Kelly ML, Herschlag D, Al-Hashimi HM. 2019. The roles of structural dynamics in the cellular functions of RNAs. Nat Rev Mol Cell Biol 20: 474–489.

28. Geary C, Baudrey S, Jaeger L. 2008. Comprehensive features of natural and in vitro selected GNRA tetraloop-binding receptors. Nucleic Acids Res 36: 1138–1152.

29. Golden BL, Kim H, Chase E. 2005. Crystal structure of a phage Twort group I ribozyme–product complex. Nat Struct Mol Biol 12: 82–89.

30. Guttman M, Rinn JL. 2012. Modular regulatory principles of large non-coding RNAs. Nature. 482: 339–346.

31. Halvorsen M, Martin JS, Broadaway S, Laederach A. 2010. Disease-Associated Mutations That Alter the RNA Structural Ensemble. PLOS Genet 6: e1001074.

32. Hartwell LH, Hopfield JJ, Leibler S, Murray AW. 1999. From molecular to modular cell biology. Nature 402: C47–C52.

33. Hayden EJ, Bendixsen DP, Wagner A. 2015. Intramolecular phenotypic capacitance in a modular RNA molecule. Proc Natl Acad Sci 112: 12444–12449.

34. Hendrix DK, Brenner SE, Holbrook SR. 2005. RNA structural motifs: building blocks of a modular biomolecule. Q Rev Biophys 38: 221–243.

35. Herschlag D. 1995. RNA Chaperones and the RNA Folding Problem (∗). J Biol Chem 270: 20871–20874.

36. Herschlag D, Bonilla S, Bisaria N. 2018. The Story of RNA Folding, as Told in Epochs. Cold Spring Harb Perspect Biol 10: a032433.

37. Huang L, Lilley DMJ. 2018. The kink-turn in the structural biology of RNA. Q Rev Biophys 51.

38. Ikawa Y, Naito D, Aono N, Shiraishi H, Inoue T. 1999. A conserved motif in group IC3 introns is a new class of GNRA receptor. Nucleic Acids Res 27: 1859–1865.

39. Ikawa Y, Nohmi K, Atsumi S, Shiraishi H, Inoue T. 2001. A Comparative Study on Two GNRA-Tetraloop Receptors: 11-nt and IC3 Motifs. J Biochem (Tokyo) 130: 251–255.

40. Ishikawa J, Fujita Y, Maeda Y, Furuta H, Ikawa Y. 2011. GNRA/receptor interacting modules: Versatile modular units for natural and artificial RNA architectures. Methods 54: 226– 238.

41. Ishikawa J, Furuta H, Ikawa Y. 2013. RNA Tectonics (tectoRNA) for RNA nanostructure design and its application in synthetic biology. WIREs RNA 4: 651–664.

42. Jaeger L, Leontis NB. 2000. Tecto-RNA: One-Dimensional Self-Assembly through Tertiary Interactions. Angew Chem Int Ed 39: 2521–2524.

43. Jaeger L, Michel F, Westhof E. 1994. Involvement of a GNRA tetraloop in long-range RNA tertiary interactions. J Mol Biol 236: 1271–1276.

44. Jaeger L, Westhof E, Leontis NB. 2001. TectoRNA: modular assembly units for the construction of RNA nano-objects. Nucleic Acids Res 29: 455–463.

45. Jumper J, Evans R, Pritzel A, Green T, Figurnov M, Ronneberger O, Tunyasuvunakool K, Bates R, Žídek A, Potapenko A, et al. 2021. Highly accurate protein structure prediction with AlphaFold. Nature 596: 583–589.

46. Klein DJ, Schmeing TM, Moore PB, Steitz TA. 2001. The kink-turn: a new RNA secondary structure motif. EMBO J 20: 4214–4221.

47. Lambert D, Leipply D, Shiman R, Draper DE. 2009. The Influence of Monovalent Cation Size on the Stability of RNA Tertiary Structures. J Mol Biol 390: 791–804.

48. Lin Z, Akin H, Rao R, Hie B, Zhu Z, Lu W, Smetanin N, Verkuil R, Kabeli O, Shmueli Y, et al. 2023. Evolutionary-scale prediction of atomic-level protein structure with a language model. Science 379: 1123–1130.

49. Mathews DH, Moss WN, Turner DH. 2010. Folding and Finding RNA Secondary Structure. Cold Spring Harb Perspect Biol 2: a003665.

50. Mauger DM, Cabral BJ, Presnyak V, Su SV, Reid DW, Goodman B, Link K, Khatwani N, Reynders J, Moore MJ, et al. 2019. mRNA structure regulates protein expression through changes in functional half-life. Proc Natl Acad Sci 116: 24075–24083.

51. Mitchell C, Polanco JA, DeWald L, Kress D, Jaeger L, Grabow WW. 2019. Responsive self-assembly of tectoRNAs with loop–receptor interactions from the tetrahydrofolate (THF) riboswitch. Nucleic Acids Res 47: 6439–6451.

52. Mládek A, Šponer JE, Kulhánek P, Lu X-J, Olson WK, Šponer J. 2012. Understanding the Sequence Preference of Recurrent RNA Building Blocks using Quantum Chemistry: The Intrastrand RNA Dinucleotide Platform. J Chem Theory Comput 8: 335–347.

53. Murphy FL, Cech TR. 1994. GAAA Tetraloop and Conserved Bulge Stabilize Tertiary Structure of a Group I Intron Domain. J Mol Biol 236: 49–63.

54. Mustoe AM, Brooks CL, Al-Hashimi HM. 2014. Hierarchy of RNA Functional Dynamics. Annu Rev Biochem 83: 441–466.

55. Phan H-D, Lai LB, Zahurancik WJ, Gopalan V. 2021. The many faces of RNA-based RNase P, an RNA-world relic. Trends Biochem Sci.

56. Pley HW, Flaherty KM, McKay DB. 1994. Model for an RNA tertiary interaction from the structure of an intermolecular complex between a GAAA tetraloop and an RNA helix. Nature 372: 111–113.

57. Ponting C, Oliver P, Reik W. 2009. Evolution and Functions of Long Noncoding RNAs. Cell 136: 629–641.

58. Roth A, Breaker RR. 2009. The Structural and Functional Diversity of Metabolite-Binding Riboswitches. Annu Rev Biochem 78: 305–334.

59. Roth A, Weinberg Z, Chen AGY, Kim PB, Ames TD, Breaker RR. 2014. A widespread self-cleaving ribozyme class is revealed by bioinformatics. Nat Chem Biol 10: 56–60.

60. Rozhdestvensky TS, Tang TH, Tchirkova IV, Brosius J, Bachellerie J, Hüttenhofer A. 2003. Binding of L7Ae protein to the K-turn of archaeal snoRNAs: a shared RNA binding motif for C/D and H/ACA box snoRNAs in Archaea. Nucleic Acids Res 31: 869–877.

61. Russell R, Zhuang X, Babcock HP, Millett IS, Doniach S, Chu S, Herschlag D. 2002. Exploring the folding landscape of a structured RNA. Proc Natl Acad Sci 99: 155–160.

62. Schneider B, Sweeney BA, Bateman A, Cerny J, Zok T, Szachniuk M. 2023. When will RNA get its AlphaFold moment? Nucleic Acids Res 51: 9522–9532.

63. She R, Chakravarty AK, Layton CJ, Chircus LM, Andreasson JOL, Damaraju N, McMahon PL, Buenrostro JD, Jarosz DF, Greenleaf WJ. 2017. Comprehensive and quantitative mapping of RNA–protein interactions across a transcribed eukaryotic genome. Proc Natl Acad Sci 114: 3619–3624.

64. Shi H, Rangadurai A, Abou Assi H, Roy R, Case DA, Herschlag D, Yesselman JD, Al-Hashimi HM. 2020. Rapid and accurate determination of atomistic RNA dynamic ensemble models using NMR and structure prediction. Nat Commun 11: 5531.

65. Shin JH, Bonilla SL, Denny SK, Greenleaf WJ, Herschlag D. 2023. Dissecting the energetic architecture within an RNA tertiary structural motif via high-throughput thermodynamic measurements. Proc Natl Acad Sci 120: e2220485120.

66. Silverman SK, Zheng M, Wu M, Tinoco I, Cech TR. 1999. Quantifying the energetic interplay of RNA tertiary and secondary structure interactions. RNA 5: 1665–1674.

67. Smith KD, Lipchock SV, Ames TD, Wang J, Breaker RR, Strobel SA. 2009. Structural basis of ligand binding by a c-di-GMP riboswitch. Nat Struct Mol Biol 16: 1218–1223.

68. Smith KD, Lipchock SV, Livingston AL, Shanahan CA, Strobel SA. 2010. Structural and Biochemical Determinants of Ligand Binding by the c-di-GMP Riboswitch,. Biochemistry 49: 7351–7359.

69. Strobel SA, Cochrane JC. 2007. RNA catalysis: ribozymes, ribosomes, and riboswitches. Curr Opin Chem Biol 11: 636–643.

70. Sudarsan N, Lee ER, Weinberg Z, Moy RH, Kim JN, Link KH, Breaker RR. 2008. Riboswitches in Eubacteria Sense the Second Messenger Cyclic Di-GMP. Science 321: 411–413.

71. Turner DH, Sugimoto N, Freier SM. 1988. RNA Structure Prediction. Annu Rev Biophys 17: 167–192.

72. Vicens Q, Cech TR. 2006. Atomic level architecture of group I introns revealed. Trends Biochem Sci 31: 41–51.

73. Weinberg Z, Barrick JE, Yao Z, Roth A, Kim JN, Gore J, Wang JX, Lee ER, Block KF, Sudarsan N, et al. 2007. Identification of 22 candidate structured RNAs in bacteria using the CMfinder comparative genomics pipeline. Nucleic Acids Res 35: 4809–4819.

74. Westhof E, Masquida B, Jaeger L. 1996. RNA tectonics: towards RNA design. Fold Des 1: R78–R88.

75. Yesselman JD, Denny SK, Bisaria N, Herschlag D, Greenleaf WJ, Das R. 2019a. Sequence-dependent RNA helix conformational preferences predictably impact tertiary structure formation. Proc Natl Acad Sci 116: 16847–16855.

76. Yesselman JD, Eiler D, Carlson ED, Gotrik MR, d’Aquino AE, Ooms AN, Kladwang W, Carlson PD, Shi X, Costantino DA, et al. 2019b. Computational design of three-dimensional RNA structure and function. Nat Nanotechnol 14: 866–873.

77. Zakrevsky P, Calkins E, Kao Y-L, Singh G, Keleshian VL, Baudrey S, Jaeger L. 2021. In vitro selected GUAA tetraloop-binding receptors with structural plasticity and evolvability towards natural RNA structural modules. Nucleic Acids Res 49: 2289–2305.

78. Zappulla DC, Cech TR. 2004. Yeast telomerase RNA: A flexible scaffold for protein subunits.

79. Proc Natl Acad Sci **101**: 10024–10029.

80. Zhang L, Doudna JA. 2002. Structural Insights into Group II Intron Catalysis and Branch-Site Selection. Science 295: 2084–2088.

81. Zhang M, Perelson AS, Tung C-S. 2011. RNA Structural Motifs. In *eLS*, John Wiley & Sons, Ltd

