## Supplemental Methods, Tables, Figures for "Exploring the energetic and conformational properties of the sequence space connecting naturally occurring RNA tetraloop receptor motifs"

*Library generation and sequencing*

The DNA library was prepared as previously described (Buenrostro et al. 2014; She et al. 2017; Denny et al. 2018; Becker et al. 2019; Jarmoskaite et al. 2019; Yesselman et al. 2019; Bonilla et al. 2021). We designed a library of 880 tectoRNA sequences 115 nucleotides in length; sequences were padded with a polyT repeat on the 5' end to maintain a consistent length. This library was flanked by common primers for PCR amplification (Oligopool_Left and Oligopool_Right, Table S1). This library was pooled with sequences for other experiments (n = 29,837 total; not discussed here) and ordered from GensScript as a 90K Custom Oligo Pool (GenScript Biotech Corporation). The library was ordered in triplicate to ensure coverage of all sequences. We quantified initial percentage of full-length library constructs (0.11 - 0.25%, or 15,000 – 33,000 library copies in 1 µl) by performing qPCR with a LUNA qPCR kit (NEB M3003); phiX genomic DNA (Illumina FC-110-3001) was used as a concentration standard. The 10 µL qPCR reactions contained 0.5 µM primers (Oligopool_Left and Oligopool_Right) and 1 µL of the library or phiX standard serially diluted in 0.1% Tween20. The library concentration was determined to be 85 pg/µL by comparing CT values against the concentration standard.

We performed one round of PCR amplification to purify the full-length sequences from the oligo pool using a Phusion High-Fidelity Polymerase (NEB M0530) with the aforementioned common primers (Oligopool_Left and Oligopool_Right) in the presence of 3% DMSO. The oligo library (1 µL) was used as the PCR template. PCR was performed with 9 cycles (3 min initial denaturation at 98 °C; cycles consisting of 30 s at 98 °C, 30 s at 62 °C, 30 s at 72 °C; 2 min final elongation at 72 °C) to limit amplification bias; the amplicons were purified using a QIAquick PCR Purification Kit (Qiagen 28104).

A five-piece PCR assembly reaction was then performed with the amplified library to append sequences required for Illumina sequencing, sequences required during the array experiment, and a 16 nt random sequence to serve as a unique barcode for each DNA molecule. The reaction contained 1 μL of the library, 137 nM each of the two outside primers (Oligo_C and Oligo_D, Table S1), and 3.84 nM each of the two adaptor oligos containing the appended sequences (C1_R1_BC_RNAP and Dark_Read2, Table S1). The PCR reaction was performed using a Phusion High-Fidelity Polymerase (with 3% DMSO) for 14 cycles of amplification (3 min initial denaturation at 98 °C; cycles consisting of 30 s at 98 °C, 30 s at 62 °C, 60 s at 72 °C; 2 min final elongation at 72 °C). The assembled library was purified using a QIAquick PCR purification kit (Qiagen 28104).

We next performed bottlenecking on the assembled sequencing library to reduce the number of unique sequences to ~650,000; this bottlenecking procedure ensures adequate coverage of each barcoded sequence for the sequencing run and subsequent array experiment.(Buenrostro et al. 2014) To do so, we quantified the assembled library by performing qPCR as described above. The library concentration was determined and a volume corresponding to 650,000 molecules was used in a final PCR reaction to prepare for sequencing. This last PCR reaction was performed using a NEBNext kit (NEB M0541) with 1.25 μM primer concentrations (Oligo_C and Oligo_D) for 25 cycles of amplification (3 min initial denaturation at 98 °C; cycles consisting of 30 s at 98 °C, 30 s at 65 °C, 30 s at 72 °C; 2 min final elongation at 72 °C). The amplified, bottlenecked library was purified using a QIAquick PCR purification kit and quantified via qPCR as described above.

The final library was pooled with fiducial (Fiducial_Chip, Table S1) and phiX (IDT) sequences for a final composition of 10% library, 1% fiducial, 89% phiX. The pooled sequences were sequenced using an Illumina MiSeq 150 v3 sequencing kit run for three cycles: 76 bases in read 1, 76 bases in read 2, and 8 bases for an i7 index read. The pooled sequence concentration was adjusted (15 pM final concentration) to yield a cluster density of ~800k clusters/mm$^2$. The sequencing output consisted of a clusterID, read 1 sequence, and read 2 sequence for every cluster on the chip. The clusterID contained information regarding the physical location of each cluster, which is used downstream to match clusters to library variants. The read 1 sequence contained the barcode sequences for each cluster, and the read 2 sequence contained the library sequence for each cluster. Clusters with common barcodes were processed to determine the consensus read 2 sequence per barcode; barcodes corresponding to poor read 2 consensus sequences or poor representation on the chip were filtered out as previously described (Denny et al. 2018; Yesselman et al. 2019). The barcodes passing quality filters were assigned to library variants by matching the reverse complement of the read 2 sequence to the library variants (850 of the 880 total library variants). Sequencing data from 4 separate MiSeq chips were pooled to map the bottlenecked barcodes to corresponding constructs.

*Flow piece preparation*

The fluorescent tectoRNA flow piece was prepared as follows. Flow piece hairpins with either a GAAA or GUAA tetraloop (Mut2_GAAA and Mut2_GUAA, Table S2) were ordered from

Integrated DNA Technologies as 5'-amino, C6-modified, HPLC-purified RNAs (IDT). The flow piece RNAs were fluorescently labelled with Cy3b-NHS and purified *via* denaturing Urea-PAGE gel electrophoresis (20% acrylamide, 8 M urea, and 1x TBE, pH 7.4 [89 mM Tris-HCl, 89 mM boric acid, and 2 mM sodium EDTA]) following established protocols (Greenfeld and Herschlag 2013; Petrov et al. 2013). The band containing purified RNA was excised from the gel and eluted into water using the crush-and-soak method. The RNA molecules were isolated from the gel fragments using a CoStar Spin-X Centrifuge Tube Filter, 0.22 μm pore size (Corning 8160); the permeate was then transferred to an Amicon Ultra-0.5 Centrifugal Filter, 3 kDa MWCO (Millipore Sigma UFC5003) and cleaned up by spinning with water. The supernatant containing purified flow piece RNA was saved, and its concentration was determined using a Qubit High Sensitivity RNA Assay kit (ThermoFischer Q32852).

*Generation of RNA on chip*

We transcribed RNA molecules on the MiSeq chip as previously described (Buenrostro et al. 2014; She et al. 2017; Denny et al. 2018; Becker et al. 2019; Jarmoskaite et al. 2019; Yesselman et al. 2019; Bonilla et al. 2021). We used a modified Illumina Genome Analyzer IIx (Illumina GAIIx) with custom temperature and fluidics control to prepare the RNA library. The MiSeq sequencing chip was processed and used to generate dsDNA and RNA as follows. The chip was incubated with 100% formamide to remove non-covalently attached molecules from the chip and treated with Cleavage buffer (100 mM Tris-HCl, 125 mM NaCl, 0.05% Tween20, 100 mM TCEP, pH 7.4) to remove fluorescent markers from the sequencing reaction.

The remaining ssDNA covalently attached to the chip was converted to dsDNA through the binding of a biotinylated primer (Biotin_Dark_Read2, Table S2) in Hybridization buffer (5x SSC [ThermoFisher 15557044], 5 mM EDTA, 0.01% Tween20). At this step, we also introduced a fluorescently labelled oligo to bind to the fiducial markers also present on the chip (Fiducial_Flow, Table S2) that was used downstream for cluster registration (mapping imaged clusters to the sequencing results). The chip was washed in Annealing buffer (1x SSC [ThermoFisher 15557044], 5 mM Na EDTA, 0.01% Tween20) followed by wash buffer (10 mM Tris HCl pH 8.0, 5 mM Na EDTA, 0.05% Tween20). dsDNA was generated using Klenow fragment (NEB M0212) in Klenow buffer (1x NEB buffer 2 [NEB B7002S], 250 μM each dNTP mix [ThermoFisher 18427013], 0.01% Tween20). To block any remaining ssDNA that was not converted to dsDNA, we hybridized a non-fluorescent oligo (Dark_Stall, Table S2) in Hybridization buffer followed by a wash with Annealing buffer. To ensure no unblocked ssDNA remained on the chip (which would bind fluorescent oligos used for RNA quantification at a later step), we flowed on a fluorescently-labelled oligo complementary to the RNA polymerase stall sequence (Fluorescent_Stall, Table S2) for imaging downstream.

Before RNA generation on the chip, we flowed on streptavidin to bind the biotinylated oligos previously used to prime dsDNA generation. Streptavidin stalls RNA polymerase so that the RNA transcript remains tethered to the chip. Next, we flowed on free biotin to saturate remaining binding sites on the bound streptavidin molecules. At this point, the chip was imaged to ensure that no Fluorescent_Stall oligos had bound and to align the locations of the fluorescently labelled fiducial clusters with their positions recorded in their clusterIDs.

RNA generation proceeded in three steps: first, the chip was incubated in Stall buffer containing 2.5 μM each of GTP, ATP, and UTP in Transcription buffer (20 mM Tris-HCl pH 8, 7 mM $MgCl_2$, 20 mM NaCl, 0.1 mM Na EDTA, 1% glycerol, 1.5 mg/mL BSA, 0.02% Tween20, 0.01 mM DTT). Transcription initiation was performed by the addition of *E. coli* RNA polymerase saturated with sigma factor 70 (NEB M0551S) in Stall buffer. The lack of CTP in the Stall buffer prevented the RNAP from continuing transcription upon reaching the first “C” in the nascent transcript and ensured that only one RNAP would be bound to each dsDNA molecule on the chip. Excess RNAP was washed away with Stall buffer. The transcription of full-length RNA was performed by the addition of Extension buffer, which contained 1 mM each NTP in Transcription buffer and 0.5 μM each of the Fluorescent_Stall oligo and Dark_Read2 oligo, which bind to the nascent RNA transcript on either side of the chip piece hairpin. The chip was then imaged to measure the Fluorescent_Stall fluorescence, which quantifies the amount of RNA generated in each cluster.

*tectoRNA equilibrium binding experiments*

Binding energies ($\Delta G_{\mathrm{bind}}$) for each of the chip piece RNAs tethered to the chip binding to a common fluorescently labelled flow piece RNA were obtained by incubating the flow piece at different concentrations and measuring the change in flow piece fluorescence at each cluster of chip piece RNAs. Flow piece RNAs dilutions were prepared in Binding buffer (89 mM Tris-Borate, pH 8.0, 0.01 mg/ml yeast tRNA, 0.01% Tween20, 30 mM $MgCl_2$, and either 150 mM or 0 mM KCl; see main text). The RNA was subject to refolding prior to flowing onto the chip (1 min 45 s at 95 °C followed by incubating on ice for 2 min; other experiments suggest that the refolding step need not be performed for each dilution and that refolding can be performed in water prior to preparation in Binding buffer and serial dilution) (Denny et al. 2018; Yesselman et al. 2019; Bonilla et al. 2021). The concentrations used across the experiments varied and are reported in Table S3. Binding experiments were performed at 22 °C, and incubations were performed for sufficient time to reach equilibrium as previously determined (Denny et al. 2018; Yesselman et al. 2019; Bonilla et al. 2021). After each equilibrium, the chip was imaged for fluorescence in the green and red channels to quantify the amount of flow piece and chip piece RNA present in each cluster, respectively. Five experiments were performed in total (Table S3).

*Image processing and cluster fitting*

The clusters present on the images were assigned to sequenced clusters by matching the position on the chip to the position information present in the clusterID, a process known as "registration," as previously described (Buenrostro et al. 2014; She et al. 2017). In brief, the cluster images were subject to an iterative cross correlation analysis with sub-pixel resolution wherein cluster densities were fit to a 2D Gaussian. Since there was little binding of the fluorescently labelled flow piece at low concentrations, we used the fluorescence of the Fiducial_Flow oligo bound to fiducial clusters to register the green channel images; we used the library clusters to register the red channel images since each library cluster will be bound by Fluorescent_Stall oligos. The cluster densities were then fit to 2D Gaussians and corrected for nonuniform fluorescence across the image, and the total integrated fluorescence values were calculated as $2\pi A\sigma^2$, where $A$ and $\sigma$ are the amplitude and standard deviation of the Gaussians, respectively. The fluorescence of the flow piece (green channel) was normalized by the fluorescence of the total RNA in each cluster (red channel) to account for variability in cluster sizes across the chip.

The normalized fluorescence data for each cluster was fit to an equilibrium binding isotherm:

$$f(x) = f_{min} + f_{max} \frac{x}{x + \exp\left(\frac{\Delta G}{RT}\right)}$$

where $f(x)$ is the normalized fluorescence as function of flow piece concentration, $f_{min}$, $f_{max}$, and $\Delta G$ are free parameters, $x$ is the flow piece concentration, $R$ is the ideal gas constant, and $T$ is temperature in Kelvin. As described previously, we pooled the $f_{min}$ from clusters corresponding to weakly-binding library variants that do not saturate flow piece binding under assayed conditions to estimate a global $f_{min}$ value and used the $f_{max}$ values from clusters corresponding to tightly-binding library variants that saturate flow piece binding to estimate the distribution for $f_{max}$ (Denny et al. 2018). Finally, we resampled the $f(x)$ values for each library variant (*e.g.*, resampled the normalized cluster fluorescence) and sampled the estimated $f_{max}$ distribution to repeatedly fit $\Delta G$ values10,000 times for each library variant to provide a bootstrapped distribution for $\Delta G$ values. As stated in the main *Methods*, library variant with fewer than 5 clusters expressed on the chip were omitted from analysis, and an upper limit for quantitative $\Delta G$ values was established per experimental condition based on library controls (Denny et al. 2018; Yesselman et al. 2019; Bonilla et al. 2021).

**Supplemental Tables and Figures**

*Table S1: Oligos used for library generation and sequencing preparation.*

| Name | Sequence |
|---|---|
| Oligopool_Left | TTGTATGGAAGACGTTCCTGGAT |
| Oligopool_Right | GCTGAACCGCTCTTCCGATCT |
| Oligo_C | AATGATACGGCGACCACCGA |
| Oligo_D | CAAGCAGAAGACGGCATACGA |
| C1_R1_BC_RNAP | AATGATACGGCGACCACCGAGATCTACACTCTTTCCCTACACGAC GCTCTTCCGATCTNNNNNNNNNNNNNNNNNTTTATGCTATAATTATT TCATGTAGTAAGGAGGTTGTATGGAAGACGTTCCTGGAT |
| Dark_Read2 | CGGTCTCGGCATTCCTGCTGAACCGCTCTTCCGATCT |
| Fiducial_Chip | AATGATACGGCGACCACCGAGATCTACACTCTTTCCCTACACGAC GCTCTTCCGATCTCTTGGGTCCACAGGACACTCGTTGCTTTCCAG ATCGGAAGAGCGGTTCAGCAGGAATGCCGAGACCGATCTCGTAT GCCGTCTTCTGCTTG |

*Table S2: Oligos used for RNA array experiments.*

RNA nucleotides are prefixed “r” and modifications correspond to IDT identifiers

| Name | Sequence |
|---|---|
| Mut2_GAAA | /5AmMC6/rCrUrArGrGrArArUrCrUrGrGrCrCrCrArUrArGrArArGrGrArArAr CrUrUrCrUrArUrGrGrGrCrCrUrGrUrGrUrCrCrUrArG |
| Mut2_GUAA | /5AmMC6/rCrUrArGrGrArArUrCrUrGrGrCrCrCrArUrArGrArArGrGrUrArAr CrUrUrCrUrArUrGrGrGrCrCrUrGrUrGrUrCrCrUrArG |
| Biotin_Dark_Read2 | /5Biosg/CAAGCAGAAGACGGCATACGAGATCGGTCTCGGCATTCCTG CTGAACCGCTCTTCCGATCT |
| Fiducial_Flow | /5TYE563/GGAAAGCAACGAGTGTCCTGTGGACCCAAG |
| Dark_Stall | GGATCCAGGAACGTCTTCCATACAACCTCCTTACTACAT |
| Fluorescent_Stall | GGATCCAGGAACGTCTTCCATACAACCTCCTTACTACAT/3AlexF546N/ |

*Table S3: Flow piece concentrations used in each experiment*

| | Experiment | | | C1 (nM) | C2 (nM) | C3 (nM) | C4 (nM) | C5 (nM) | C6 (nM) | C7 (nM) | C8 (nM) |
|---|---|---|---|---|---|---|---|---|---|---|---|
| | TL | $MgCl_2$ (mM) | KCl (mM) | | | | | | | | |
| 1 | GAAA | 30 | 0 | 0.74 | 2.24 | 6.7 | 20 | 61 | 181 | 544 | 1634 |
| 2 | GAAA | 30 | 0 | 0.91 | 2.7 | 8.2 | 25 | 74 | 222 | 666 | 2000 |
| 3* | GAAA | 30 | 150 | 9.1 | 27 | 1 | 3 | 82 | 222 | 666 | 2000 |
| 4 | GUAA | 30 | 0 | 0.91 | 2.7 | 8.2 | 25 | 74 | 222 | 666 | 2000 |
| 5 | GUAA | 30 | 150 | 0.91 | 2.7 | 8.2 | 25 | 74 | 222 | 666 | 2000 |

* Following the C2 measurement, flow piece RNA was washed off with Binding buffer and the chip was imaged to ensure no flow piece remained bound prior to the C3 measurement.

*Dataset S1 (separate file): Binding data for the TLR variants*

The dataset contains $\Delta G_{bind}$ values for the 44 TLR variants binding to a GAAA or GUAA tetraloop in the presence or absence of 150 mM $K^+$. Data represent average $\Delta G_{bind}$ values over a subset of the tectoRNA scaffolds; the median and 95% confidence intervals are reported (calculated as described in the *Methods*). Variants which resulted in limits to binding are denoted as such, and their confidence intervals therefore lack an upper bound.

*Dataset S2 (separate file): Thermodynamic fingerprint data for the TLR variants*

The dataset contains $\Delta\Delta G_{rel}$ values for TLR pathway intermediates compared to their "wild type" receptors (11ntR and IC3R or Vc2R). Data for 11ntR → IC3R pathway variants are presented in Sheet 1, "IC3R_Pathway_Fingerprints", with median and 95% confidence intervals for $\Delta\Delta G_{rel}$ and the adjusted *p*-value testing $\Delta G_{rel} \neq 0$ reported for each scaffold. Likewise, data for 11ntR → Vc2R pathway variants are presented in Sheet 2, "Vc2R_Pathway_Fingerprints".

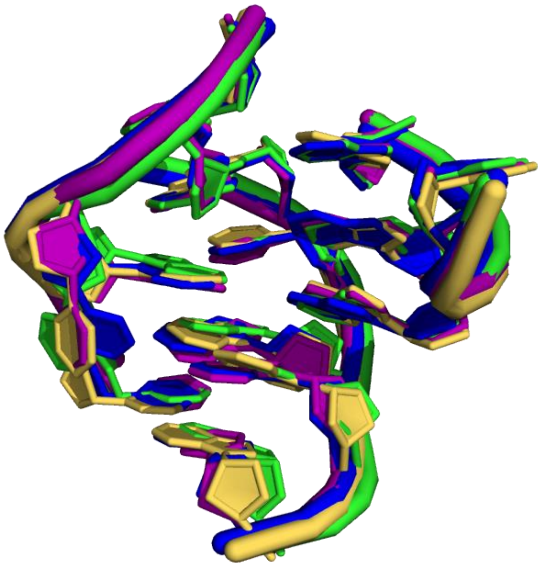

*Figure S1: 11ntR sequences from different host RNAs have the same 3D structure*

The 3D structures of the 11ntR and its variants found in natural RNAs: 11ntR in P4P6 from *Tetrahymena* Group I intron [blue, PDB: 1GID (Cate et al. 1996)], G6A variant found in Group II intron [yellow, PDB: 4R0D (Robart et al. 2014)], G6C variant found in Group II intron [pink, PDB: 6ME0 (Haack et al. 2019)], A4C variant in RNase P [green, PDB: 1NBS (Krasilnikov et al. 2003)]. The structures align with <1 Å RMSD, with the largest variation occurring in the flipped out U9 residue.

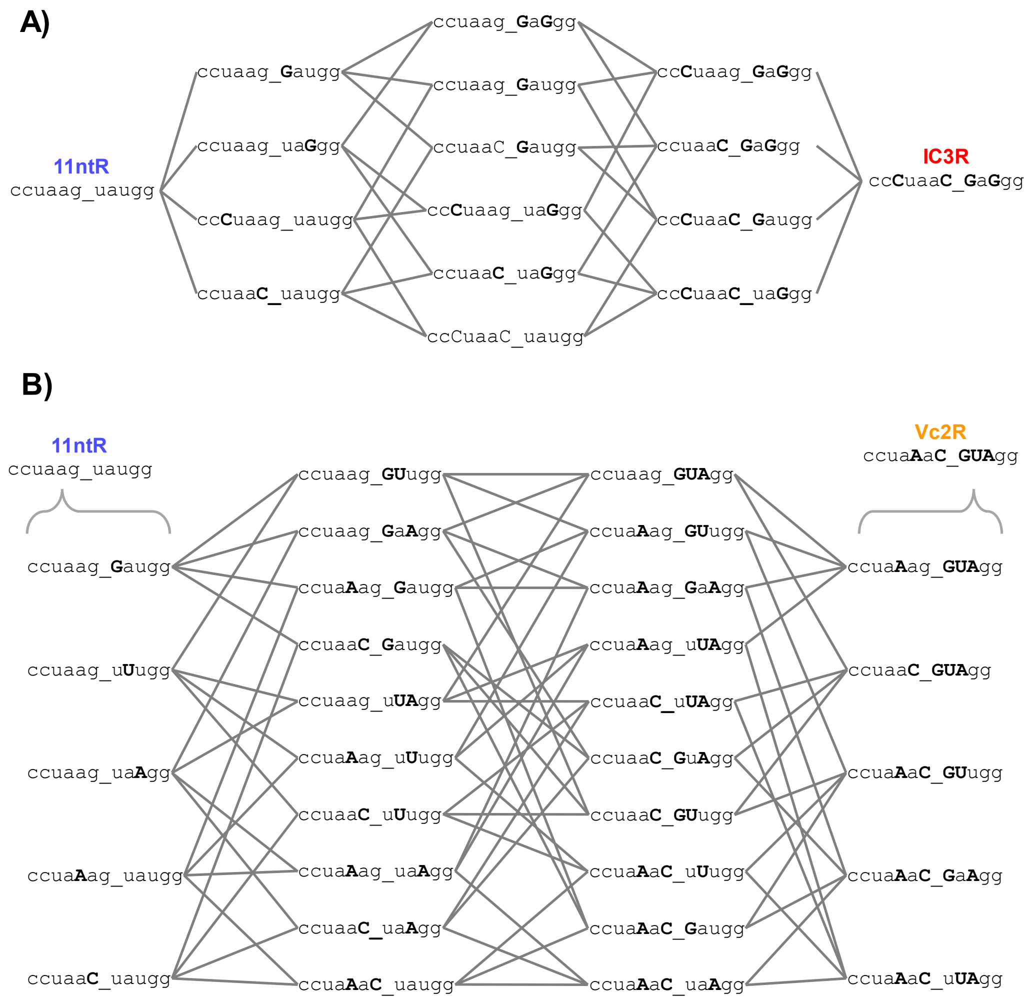

*Figure S2: Mutation pathways from the 11ntR to the IC3R and Vc2R*

The intermediate TLR sequences that comprise the mutational pathways from the 11ntR to the IC3R (A) and the Vc2R (B). Lowercase letters represent the wild type 11ntR sequence, and bolded uppercase letters indicate mutated residues from the 11ntR. Residues are ordered as in the numbering introduced in Figure 1, e.g., the leftmost residue is position “1” and the final residue is position “11”. Grey lines connect TLR intermediates separated by a single mutation.

```
 8.1      8.2      8.3      8.4      8.5      9.1      9.2      9.3      9.4      9.5
 G A      G A      G A      G A      G A      G A      G A      G A      G A      G A
G - A    G - A    G - A    G - A    G - A    G - A    G - A    G - A    G - A    G - A
G - C    G - C    G - C    G - C    G - C    G - C    G - C    G - C    G - C    G - C
U - A    U - A    U - A    U - A    U - A    C - G    C - G    C - G    C - G    G - C
G - C    G - C    G - C    C - G    G - C    G - C    U - A    U - A    G - C    U - A
A • A    A • G    |   G    G - C    C • U    G - C    G - C    G - C    C - G    A - U
|   A    |   A    |   G    A   |    |   A    A • A    C - G    G - C    G • A    C - G
U - A    |   A    A - U    A   |    |   A    A   |    A   |    |   G    |   A    A - U
C - G    U - A    U - A    U - G    U - A    G - C    G - C    |   G    C - G    A - U
U - A    C - G    C - G    G - C    C - G    C - G    C - G    C - G    C - G    U - A
A - U    U - A    U - A    U - A    U - A    A - U    A - U    C - G    A - U    C - G
         A - U    A - U    A - U    A - U    A - U    A - U    A - U    A - U
                                                               A - U

10.1     10.2     10.3     10.4     10.5     11.1     11.2     11.3     11.4     11.5
 G A      G A      G A      G A      G A      G A      G A      G A      G A      G A
G - A    G - A    G - A    G - A    G - A    G - A    G - A    G - A    G - A    G - A
G - C    G - C    G - C    G - C    G - C    G - C    G - C    G - C    G - C    G - C
C - G    C - G    C - G    C - G    C - G    C - G    C - G    C - G    C - G    C - G
G - C    U - A    U - A    U - A    U - A    U - A    U - A    U - A    U - A    U - A
C - G    G - C    G - C    G - C    G - C    U - A    U - A    U - A    U - A    U - A
C • A    A - U    U - A    C - G    G - C    G - C    C • C    A • C    G - C    G • G
A • C    G • G    C • C    U • U    |   G    G • G    G • G    U • U    |   U    U • U
G - C    U - A    A - U    |   U    |   G    G • G    G • G    U • U    A - U    U • U
C - G    C - G    C - G    C - G    C - G    C • U    C - G    C - G    U - G    C - G
A - U    A - U    A - U    C - G    C - G    U - A    U - A    U - A    C - G    U - A
A - U    A - U    A - U    A - U    G - C    A - U    A - U    A - U    U - A    A - U
                           A - U    A - U    A - U    A - U    A - U    A - U    A - U
                                    A - U                               A - U
```

*Figure S3: Chip-piece scaffold sequences and their predicted secondary structures*

The 20 scaffold sequence variants used in this study (bold) shown with the GAAA tetraloop (unbolded). Predicted Watson-Crick base pairs are shown by solid lines and mismatches by "•". The scaffolds are sorted by the Watson-Crick + mismatched base paired lengths. The construct number corresponds to the number of opposing residues, whether Watson-Crick or non-Watson-Crick pairs; the decimal refers to the individual constructs of each length.

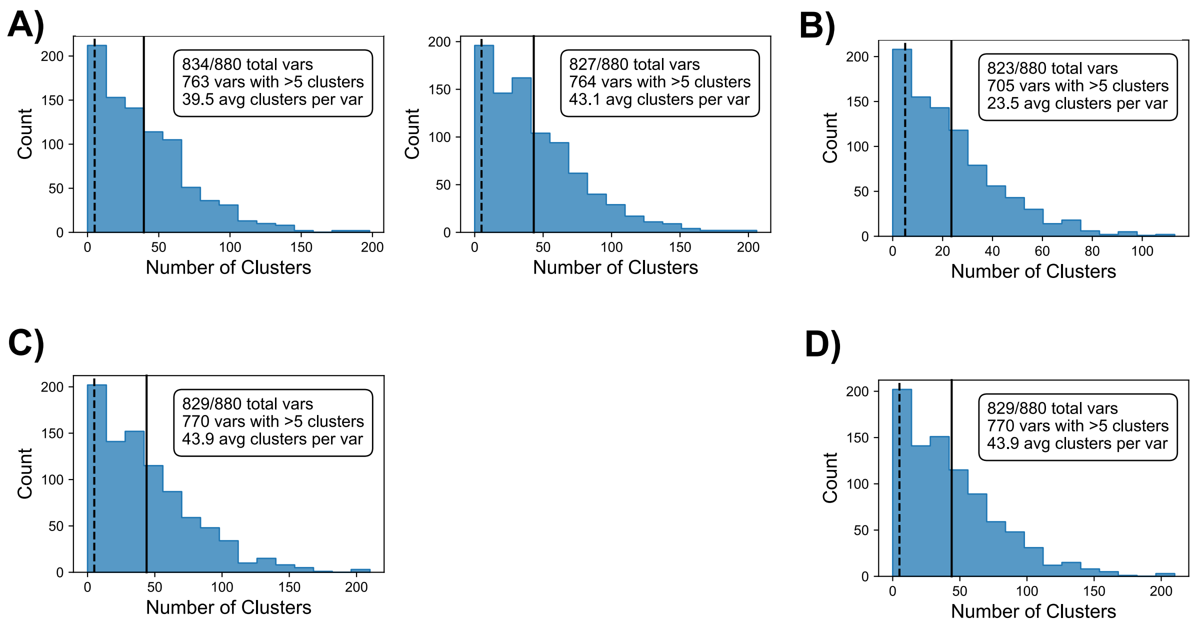

*Figure S4: The number of clusters per sequence variant present on-chip.*

Plots depict the distribution of the number of clusters per library variant present on the MiSeq chip for (A) the two experiments performed with GAAA and no $K^+$, (B) the experiment performed with GUAA and no $K^+$, (C) the experiment performed with GAAA and 150 mM $K^+$, and (D) the experiment performed with GUAA and 150 mM $K^+$. Solid vertical lines depict the average number of clusters per variant and the dashed line marks $n$ = 5 clusters per variant.

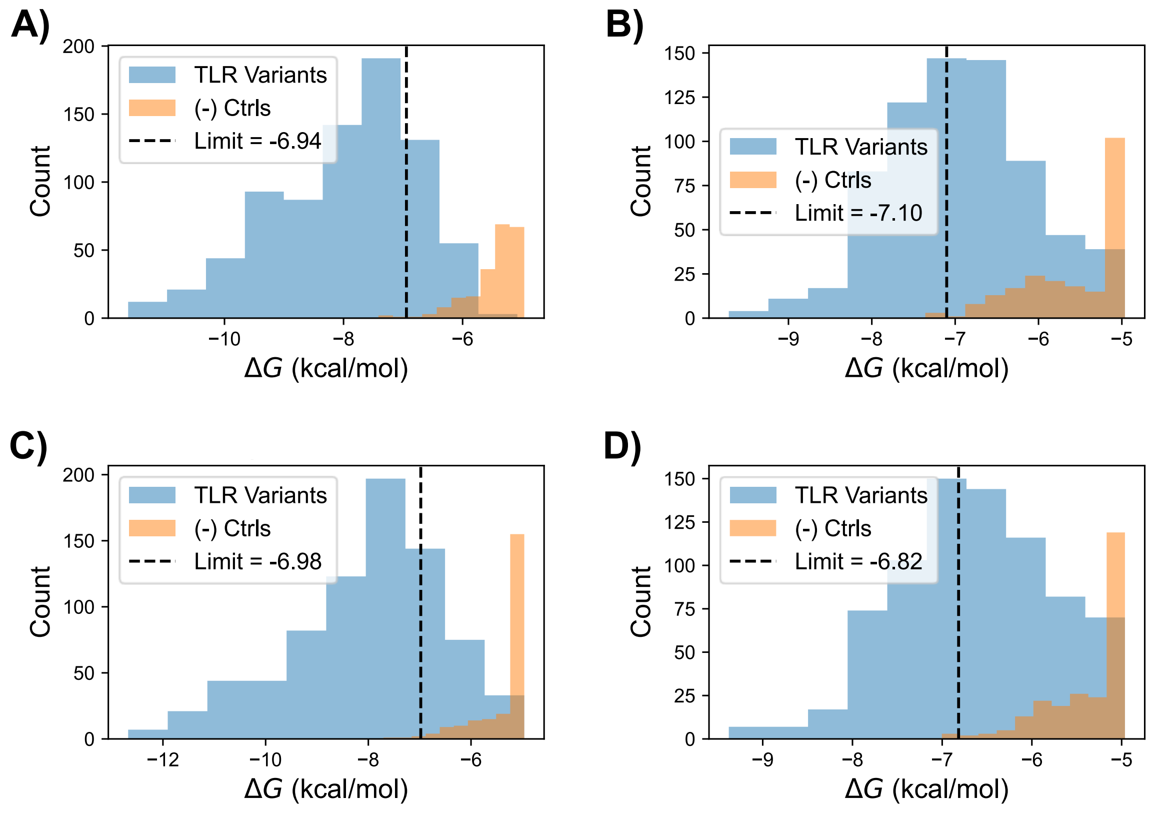

*Figure S5: Lower limit for quantitative $\Delta G_{bind}$ values*

Plots depict the distribution of $\Delta G_{\text{bind}}$ values for TLR library variants (blue) and a negative control library containing no TLR sequence (orange) for (A) the average of the two experiments performed with GAAA and no $K^+$, (B) the experiment performed with GUAA and no $K^+$, (C) the experiment performed with GAAA and 150 mM $K^+$, and (D) the experiment performed with GUAA and 150 mM $K^+$. Dashed vertical lines mark the limit for each experiment, defined as the 99$^{th}$ percentile of the negative control library affinities.

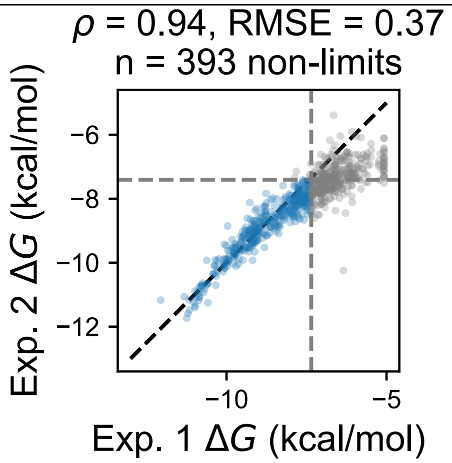

*Figure S6: Comparison of replicate experimental data*

Δ*G* values for all variants with greater than 5 clusters on the chip (n = 748) for the two experiments performed with GAAA and no $K^+$. Blue points represent library variants with quantitative (non-limit) Δ*G* values (n = 393) and grey points represents library variants that resulted in limits. Dashed gray lines represent the limits for the two experiments. The Pearson correlation coefficient and root mean squared error (RMSE) are calculated using the library variants with quantitative Δ*G* values.

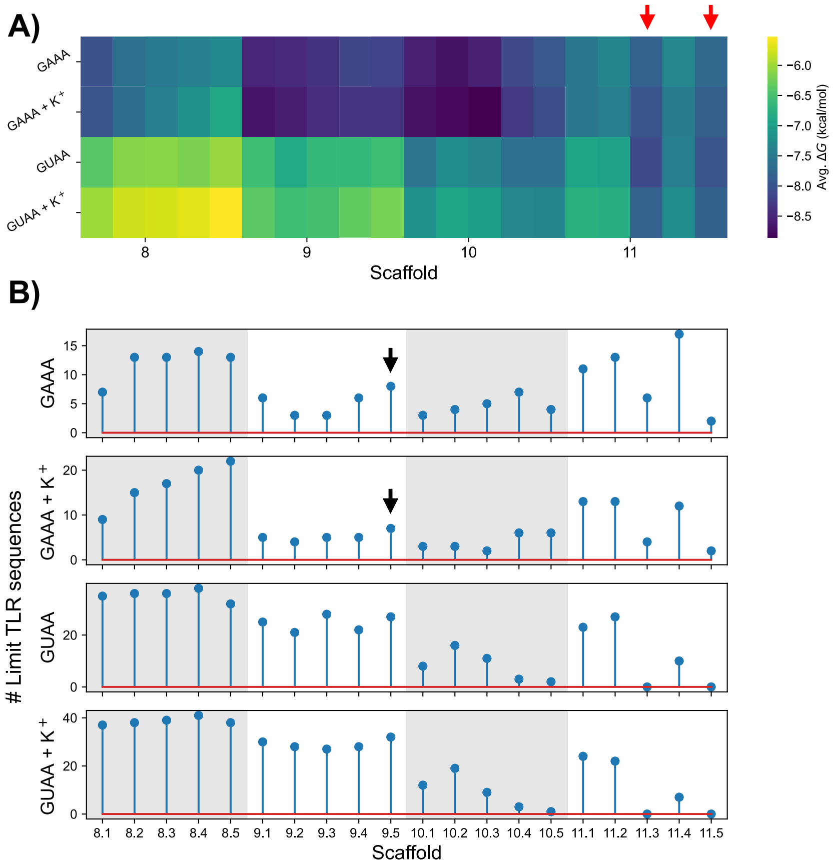

*Figure S7: Determination of the 9 scaffolds used to evaluate energy landscapes*

(A) The heatmap depicts the average $\Delta G_{\mathrm{bind}}$ for all TLR variants in a given scaffold across the four experimental conditions. TLR variants bound tighter to the 9 bp and 10 bp scaffolds than the 8 bp or 11 bp scaffolds on average, so data from 8 bp and 11 bp scaffolds were omitted when calculating $\Delta G_{\mathrm{bind}}$ measurements averaged over scaffolds. Scaffold 11.3 and 11.5 (red arrows) also resulted in tight binding, but the former resulted in conformational changes and the latter sequence was not present with the 11ntR on the array such $\Delta\Delta G$ values could not be derived. (B) The number of TLR variants with non-quantitative $\Delta G_{\mathrm{bind}}$ measurements (limits) for each scaffold under the 4 experimental conditions. Scaffold 9.5 resulted in a higher number of limit values for several conditions (black arrow) and was thus further excluded from determining average $\Delta G_{bind}$ values.

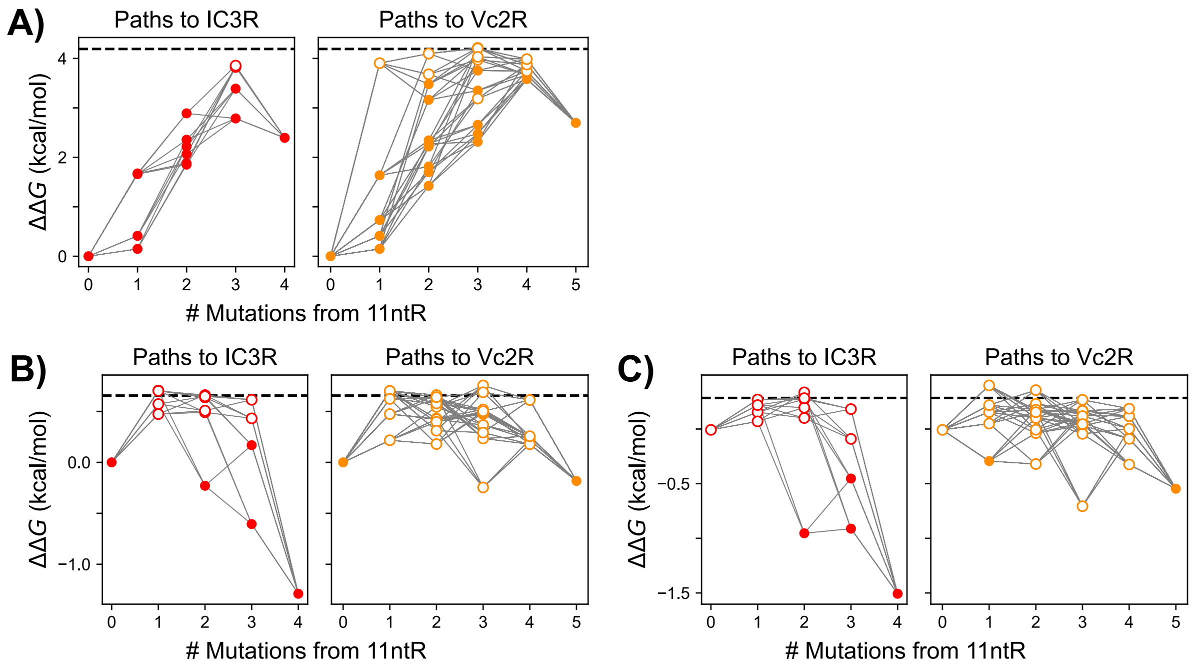

*Figure S8: Sequence-energy landscapes under various ionic and tetraloop binding conditions*

(A) The 11ntR → IC3R (red, left) and 11ntR →Vc2R (orange, right) sequence-energy landscape when binding to GAAA without $K^+$ present. ΔΔ*G*s are taken from the wild type 11ntR sequence and averaged over tectoRNA scaffolds; median values are plotted. TLRs that resulted in non-quantitative ΔΔ*G*s are represented by open points. TLR intermediates that lie on the same mutational pathways (*e.g.*, single mutants) are connected by grey lines. (B) Same as in (A) but with TLR sequences binding to GUAA in the presence of 150 mM $K^+$. (C) Same as in (B) but with TLR sequences binding to GUAA in the absence of $K^+$.

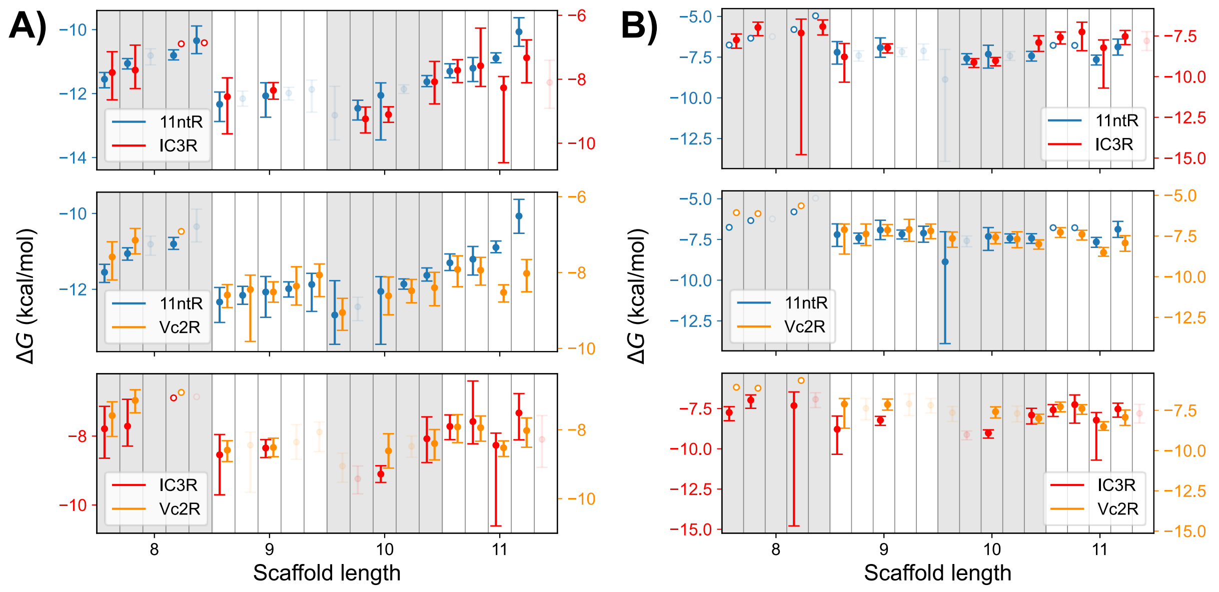

*Figure S9: Thermodynamic fingerprints wild type TLR sequences*

$\Delta G_{bind}$ values for the 11ntR (blue), IC3R (red), or Vc2R (orange) binding to (A) GAAA or (B) GUAA TL. The color of the y-axes labels correspond to the color of the TLR fingerprints, where the two y-axes are offset to facilitate comparisons of the fingerprints. Fingerprints are plotted pairwise, with the corresponding TLR sequences shown in the legend. Plotted values represent median and 95% confidence interval values; open circles represent values that are lower limits to $\Delta G_{bind}$.

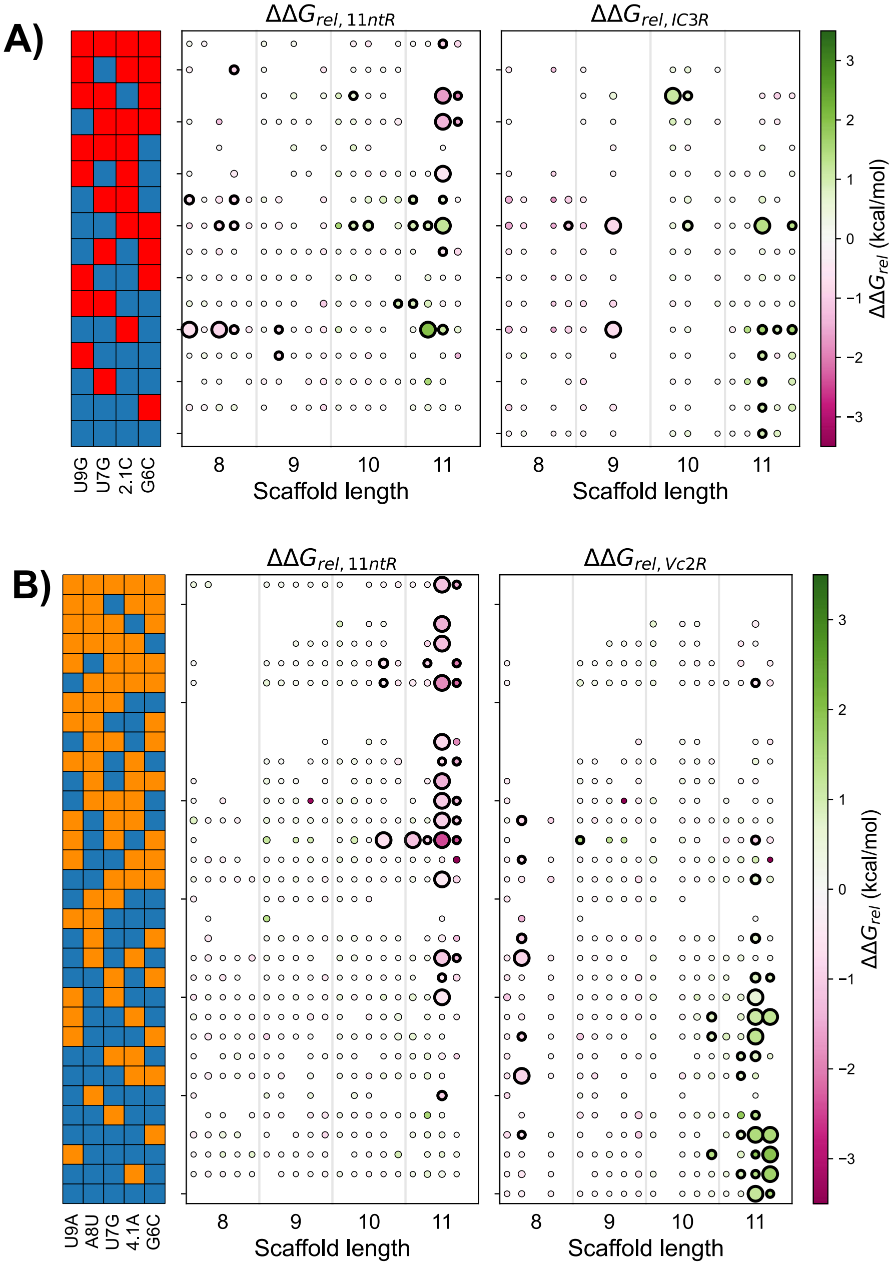

*Figure S10: Thermodynamic fingerprints for all sequence intermediates binding to the GAAA-containing flowpiece*

(A) $\Delta\Delta G_{rel}$ values for all scaffolds for TLR intermediates along the 11ntR → IC3R pathways. $\Delta\Delta G_{rel}$ values are presented relative to both the 11ntR (left) and the IC3R (right). Points represent individual $\Delta\Delta G_{rel}$ values across the 20 scaffolds, with the color corresponding to the values and larger size corresponding to greater statistical significance; $\Delta\Delta G_{rel}$ that are significantly non-zero are outlined in black. (B) Same as in (A) but for TLR intermediates along the 11ntR → Vc2R pathways. $\Delta\Delta G_{rel}$ values are calculated relative to the 11ntR (left) and the Vc2R (right).

**Supplemental References**

Becker WR, Jarmoskaite I, Vaidyanathan PP, Greenleaf WJ, Herschlag D. 2019. Demonstration of protein cooperativity mediated by RNA structure using the human protein PUM2. *RNA* **25**: 702–712.

Bonilla SL, Denny SK, Shin JH, Alvarez-Buylla A, Greenleaf WJ, Herschlag D. 2021. High-throughput dissection of the thermodynamic and conformational properties of a ubiquitous class of RNA tertiary contact motifs. *Proc Natl Acad Sci* **118**.

Buenrostro JD, Araya CL, Chircus LM, Layton CJ, Chang HY, Snyder MP, Greenleaf WJ. 2014. Quantitative analysis of RNA-protein interactions on a massively parallel array reveals biophysical and evolutionary landscapes. *Nat Biotechnol* **32**: 562–568.

Cate JH, Gooding AR, Podell E, Zhou K, Golden BL, Kundrot CE, Cech TR, Doudna JA. 1996. Crystal Structure of a Group I Ribozyme Domain: Principles of RNA Packing. *Science* **273**: 1678–1685.

Denny SK, Bisaria N, Yesselman JD, Das R, Herschlag D, Greenleaf WJ. 2018. High-Throughput Investigation of Diverse Junction Elements in RNA Tertiary Folding. *Cell* **174**: 377–390.

Greenfeld M, Herschlag D. 2013. Chapter Fifteen - Fluorescently Labeling Synthetic RNAs. In *Methods in Enzymology* (ed. J. Lorsch), Vol. 530 of *Laboratory Methods in Enzymology: RNA*, pp. 281–297, Academic Press

Haack DB, Yan X, Zhang C, Hingey J, Lyumkis D, Baker TS, Toor N. 2019. Cryo-EM Structures of a Group II Intron Reverse Splicing into DNA. *Cell* **178**: 612-623.e12.

Jarmoskaite I, Denny SK, Vaidyanathan PP, Becker WR, Andreasson JOL, Layton CJ, Kappel K, Shivashankar V, Sreenivasan R, Das R, et al. 2019. A Quantitative and Predictive Model for RNA Binding by Human Pumilio Proteins. *Mol Cell* **74**: 966–981.

Krasilnikov AS, Yang X, Pan T, Mondragón A. 2003. Crystal structure of the specificity domain of ribonuclease P. *Nature* **421**: 760–764.

Petrov A, Wu T, Puglisi EV, Puglisi JD. 2013. Chapter Seventeen - RNA Purification by Preparative Polyacrylamide Gel Electrophoresis. In *Methods in Enzymology* (ed. J. Lorsch), Vol. 530 of *Laboratory Methods in Enzymology: RNA*, pp. 315–330, Academic Press

Robart AR, Chan RT, Peters JK, Rajashankar KR, Toor N. 2014. Crystal structure of a eukaryotic group II intron lariat. *Nature* **514**: 193–197.

She R, Chakravarty AK, Layton CJ, Chircus LM, Andreasson JOL, Damaraju N, McMahon PL, Buenrostro JD, Jarosz DF, Greenleaf WJ. 2017. Comprehensive and quantitative mapping of RNA–protein interactions across a transcribed eukaryotic genome. *Proc Natl Acad Sci* **114**: 3619–3624.

Yesselman JD, Denny SK, Bisaria N, Herschlag D, Greenleaf WJ, Das R. 2019. Sequence-dependent RNA helix conformational preferences predictably impact tertiary structure formation. *Proc Natl Acad Sci* **116**: 16847–16855.
